# Old-New Recognition Memory Revisited

**DOI:** 10.1101/2025.11.11.687844

**Authors:** Xiangyu Wei, Randy L. Buckner

## Abstract

Identifying old items during recognition memory tasks elicits widespread cortical responses. Recent results suggest that a distributed network, labeled the Parietal Memory Network (PMN), responds to old items potentially reflecting mnemonic familiarity. An independent, parallel line of evidence suggests that the same network responds to salient stimuli in non-mnemonic contexts, referring to the network as the Salience Network (SAL). Here we examined responses during old-new recognition and non-mnemonic detection tasks in intensively scanned individuals to directly contrast component processes related to mnemonic familiarity with more general processes associated with target detection. In an oddball detection paradigm devoid of traditional memory demands, SAL/PMN was robustly activated. SAL/PMN was also activated by old words in a standard old-new recognition task, thus reproducing both previously reported effects. In a critical oppositional contrast, recognition was examined when the task was to detect uncommon old words versus when the task was to detect uncommon new words. SAL/PMN was activated during the detection of targets even when those targets were new items, consistent with a response to the task relevance of the targets not mnemonic familiarity. Exploratory analyses further suggested that a right-lateralized network, referred to as Frontoparietal Network B (FPN-B), responds to mnemonic familiarity. These results revise our understanding of brain networks contributing to old-new recognition separating component processes common to many forms of target detection task from processes preferentially associated with mnemonic history.

## Introduction

A classical approach to studying memory is to have participants detect old and new items in mixed lists (Galton, 1883; Strong, 1913). Such item recognition tests – also called old-new or yes-no recognition tests – engage mnemonic processes because the critical decision feature differentiating the old from new items is their history of exposure, not cues available in the immediate environment. When old items are studied extensively through multiple repetitions, recognition decisions rely primarily on familiarity – a rapidly triggered awareness that the item is old, without necessarily recalling the orifinal study episode (Mandler, 1980)^1^. The present work investigates brain networks engaged when extensively studied old items are recognized.

Old-new recognition tests are a mainstay of human electrophysiological and neuroimaging research (e.g., Paller & Kutas, 1992; Kapur et al., 1995; Nyberg et al., 1995; Rugg et al., 1996; Henson et al., 2000; Rugg & Curran, 2007), including extensive research from our own laboratory (e.g., Buckner et al., 1998; Konishi et al., 2000; Donaldson, Petersen, & Buckner, 2001; Wheeler & Buckner, 2003; Shannon & Buckner, 2004). Recent explorations of memory have frequently adopted old-new recognition paradigms (e.g., Wang et al., 2018; Gilmore, Nelson, et al., 2019; Petrovská et al., 2021; Layher et al., 2023; Koslov, Kable, & Foster, 2024; Roberts, Meade, & Fernandes, 2025). Results suggest distributed cortical and subcortical regions respond more to old than new items, including a widely reproduced parietal old / new effect (Wagner et al., 2005; Vilberg & Rugg, 2008) and varied results involving the hippocampal formation (Henson, 2005).

However, even in their simplest form, old-new recognition tests are deceptively complex tasks that involve cognitive effort, implicit effects of repetition, preparation of the response, and decision-making processes that are not specific to memory retrieval ^2^. Studies seeking to disentangle component processes supporting recognition decisions indicate that separate regions respond to distinct aspects of the decision (e.g., Dobbins et al., 2002; Wheeler and Buckner, 2003; Layher et al., 2023). Furthermore, direct contrast of old-new paradigms with other forms of memory test yield distinct brain response patterns (Chen et al., 2017).

One confounding aspect of old-new recognition decisions is that they are asymmetrical decisions^3^. In their typical implementation, responses to old items are faster. For example, in an early analysis of response times in old-new paradigms, Okada (1971) found that the speed of correct responses to new items (Correct Rejections; CR) was slower than correct responses to old items (Hits). The response time difference grew as a function of how recently the old items were studied, suggesting that the greater the strength of the mnemonic trace, the greater the response asymmetry. In classic memory theory, when the study set is large and retrieval must be based on long-term memory, the asymmetry of responding is attributed to “new” responses relying on an exhaustive search of all possibilities, whereas faster old responses arise from self-termination of the decision when a match to the memory set is detected (e.g., Ratcliff, 1978).

Paradigm variations can minimize the response time asymmetry or even reverse it. For example, decisions that require confidence ratings or remember / know judgements can change response time distributions, including in neuroimaging studies (see Figure 1 in Wheeler & Buckner, 2004). Nonetheless, under commonly investigated conditions in the field of neuroimaging, the response asymmetry in old-new recognition tests is robust. For example, the response time asymmetry can be seen in early event-related functional MRI (fMRI) studies of recognition memory (see Figure 5 of Shannon & Buckner, 2004) as well as recent studies (see Figure 2 of Chen et al., 2017). Even when conditions manipulate difficulty, CR responses within each condition are slower than to Hits (see Figure 2 of Velanova et al., 2003 and Figure 1 of Wang et al., 2018). The response asymmetry suggests that, in addition to the mnemonic history supporting differences between old and new items, the decision processes linked to detection of old items may include a form of target detection. Most broadly, these details of how participants perform memory tests are critical to understand how brain networks support mnemonic decision processes (Aminoff et al., 2015; Layher et al., 2023).

**Figure 1.**
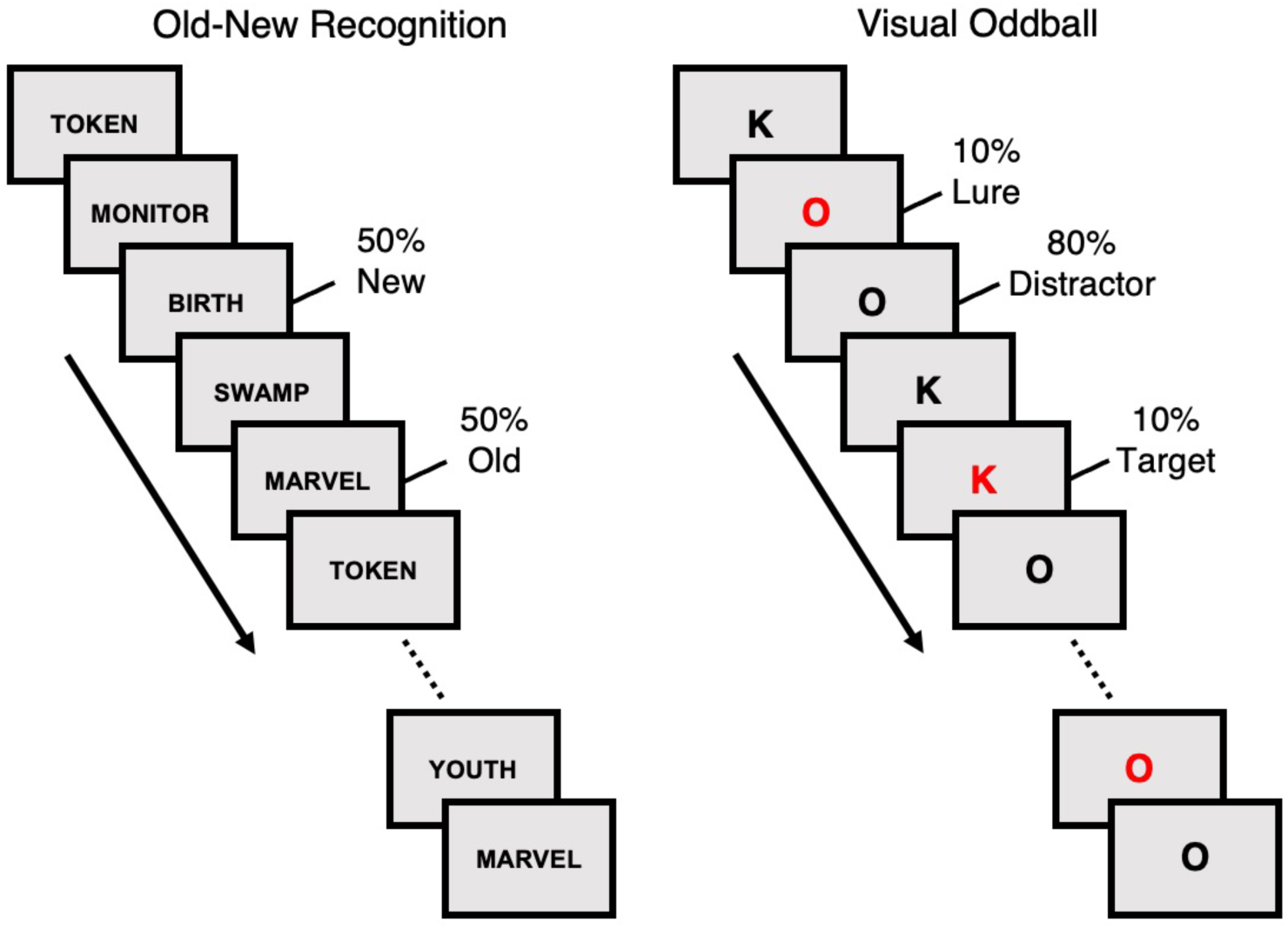
Traditional Old-New Recognition and Visual Oddball tasks. Two paradigms, arising from different experimental lineages, were examined to determine whether they activated the SAL/PMN network. (Left Panel) The Old-New Recognition task examined memory recognition under typical conditions where half of the items are old and half new. (Right Panel) The Visual Oddball task was a canonical oddball detection task. Participants responded to uncommon salient targets (red K’s) and ignored all distractors (black O’s and K’s) and lures (red O’s). The targets occurred on 10% of the trials.

**Figure 2.**
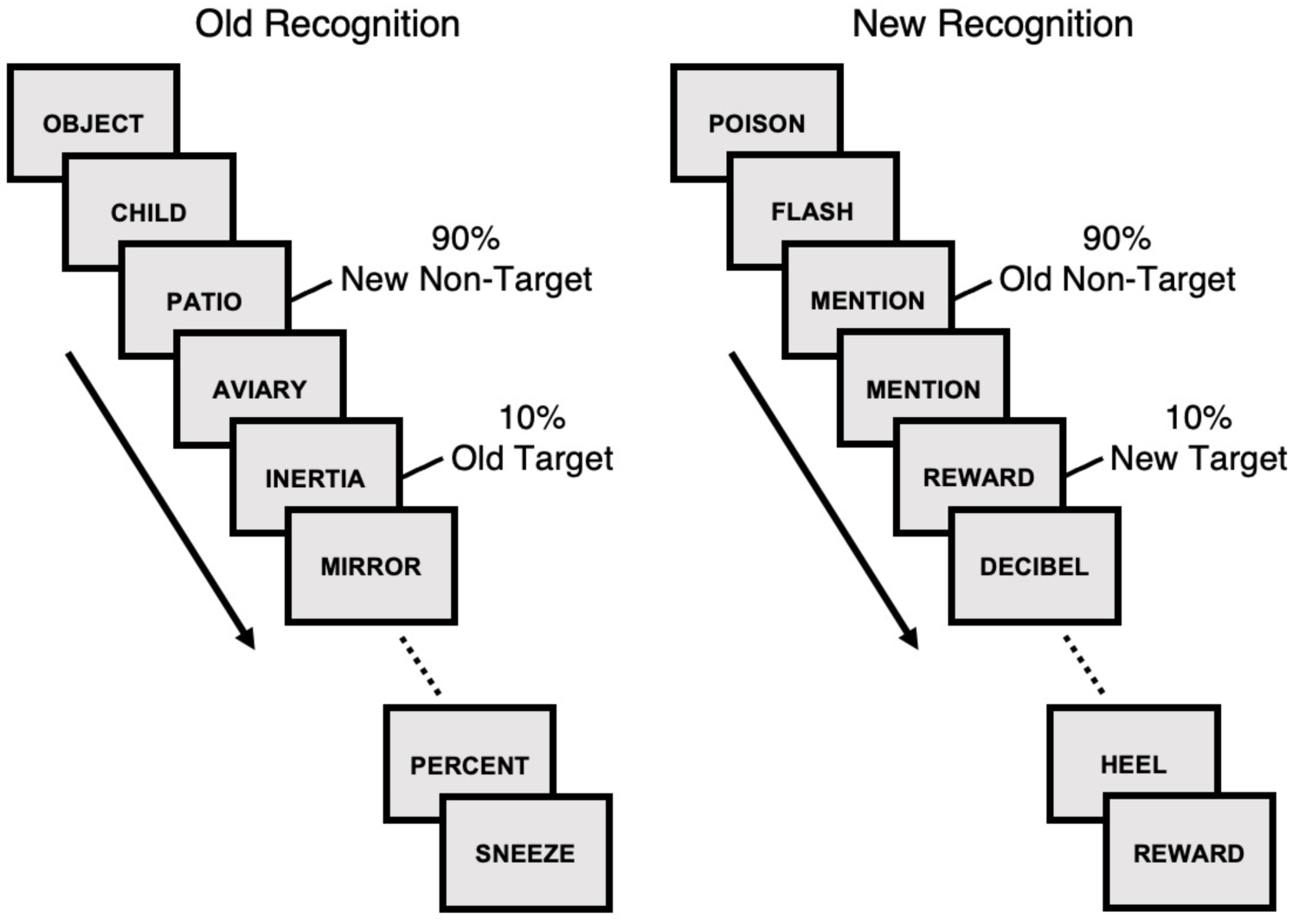
Old Recognition versus New Recognition as a hypothesis-driven oppositional contrast. Contrasting paradigms were examined that varied whether old (familiar) or new items were the target. (Left Panel) The Old Recognition task aligned the saliency and familiarity effects by designating the uncommon old words as the salient targets. (Right Panel) The New Recognition task opposed salience from familiarity by designating the new words as the salient targets. A familiarity hypothesis predicts a response to targets in the Old Recognition task but not the New Recognition task. By contrast, a salience hypothesis predicts a response to new items in the New Recognition task.

Also relevant to understanding how brain networks support recognition memory are recently developed precision neuroimaging approaches (Laumann et al., 2015; Braga & Buckner, 2017; Gordon et al., 2017; DiNicola, Braga, & Buckner, 2020; Du et al., 2024; see also Tobyne et al., 2018). Within-individual neuroimaging approaches provide opportunities to distinguish spatially adjacent networks from one another with high precision, including differentiating juxtaposed regions along the posterior midline that have been implicated in memory. Originally considered to be an integration zone (or hub) where multiple networks converge (Buckner, Andrews-Hanna, & Schacter, 2008; Andrews-Hanna et al., 2010; see also Leech, Braga, & Sharp, 2012; Braga et al., 2013), precision approaches have recently revealed the posterior midline possesses small side-by-side regions that participate in distinct distributed networks (Peer et al., 2015; Braga & Buckner, 2017; Silson et al., 2019; Braga et al., 2019; Gilmore, Nelson, et al., 2019; Kong et al., 2021; Du et al., 2024; Kwon et al., 2025). The functional properties of these distinct networks, including their midline components, has been a topic of debate.

One network that contains regions along the posterior midline is the Salience/ Parietal Memory Network (SAL/PMN), whose naming here reflects the convergence of two historical lines of research that may have targeted the same network from distinct perspectives (Du et al., 2024; see also Kwon et al., 2025; Ladwig et al., 2025). The PMN was originally identified as the network responding to repeating (familiar) items, with particular anatomical focus on the precuneus (PCU), mid-cingulate cortex (MCC), and inferior parietal lobule (IPL) (Gilmore, Nelson, & McDermott, 2015). The spatial organization of the PMN is intriguing when considered in relation to the broader memory literature because it is adjacent to (but distinct from) regions that participate in the so-called default network (DN) that is active during autobiographical remembering (Gilmore, Nelson, & McDermott, 2015; see also Du et al., 2024). In addition, the PMN is functionally coupled to the posterior hippocampus while the DN is coupled to the anterior hippocampus (Zheng et al., 2021; Angeli et al., 2025). Regions within the PMN exhibit repetition enhancement with increasing response to repeated exposure to stimuli (Gilmore, Nelson, & McDermott, 2015; Gilmore, Kalinowski, et al., 2019; Gilmore, Nelson, et al., 2019; see also McDermott et al., 2017) but are not active during autobiographical remembering (see Supplementary Figure 17 in Du et al., 2024; Angeli et al., 2025). In a recent study of old-new recognition using high-field imaging and intensive within-individual sampling, a response could be observed along PCU and MCC in many participants (Koslov, Kable, & Foster, 2024). Moreover, in group-average maps the response in the posterior midline is remarkably similar to the estimated topography of PMN (see Figure 2 of Koslov, Kable, & Foster, 2024). In a meta-analysis of old-new effects in recognition tasks, a consistent response was detected in PCU, MCC^4^, and IPL – regions implicated as components of the PMN (Kim et al., 2013). These findings suggest a role, in some manner, for the PMN in detection of mnemonic familiarity but not a broader role in autobiographical remembering.

Arising from an independent line of investigation, the SAL network refers to a group of brain regions that respond to attention-grabbing, salient stimuli, whether due to perceptual novelty, emotional intensity, or homeostatic relevance (Seeley, 2019). In a landmark study using resting-state functional connectivity, Seeley et al. (2007) described the SAL network emphasizing the dorsal anterior cingulate cortex (dACC) and the ventral anterior insula (vAI), both of which are associated with detecting and integrating salient events (Menon & Uddin, 2010; Seeley, 2019; see also Corbetta & Shulman, 2002). These regions exhibit transient responses in diverse contexts, suggesting that their roles go beyond task-specific domains to encode general salience (Critchley et al., 2004; Critchley, 2005; Bartra, McGuire, & Kable, 2013).

Despite the dual origins of their discovery, recent observations suggest that the SAL network and PMN might be the same network studied from distinct perspectives. First, detailed analyses of the functional correlation patterns reveal that the regions emphasized as components of the PMN (including PCU, MCC, and IPL) are also strongly (and selectively) correlated with the regions emphasized in studies of the SAL network (dACC and vAI) (Du et al., 2024; see also Kwon et al., 2025; Ladwig et al., 2025). Even in studies that assume the networks are separate in *a priori* models, the distributed sets of regions from both networks reveal their presence in the raw correlation structure. For example, in Lynch et al. (2024)’s analysis of the SAL network, the posterior midline was not included in the model. Nonetheless, seed-based raw correlations from dACC revealed correlations in the canonical PMN regions along the posterior midline (see Extended Data Figure 5 in Lynch et al., 2024). Second, in studies of the striatum, placing seed regions within the SAL-network-linked portion of the ventral striatum produces a distributed cortical pattern that also includes the PMN midline regions (PCU and MCC)(Kosakowski et al., 2025). Finally, the PMN network has been shown to correlate with the posterior hippocampus (Zheng et al., 2021). As would be expected based on the hypothesis that the SAL and PMN networks are the same network, the relevant portion of the posterior hippocampus responds transiently to targets in an oddball detection task but not autobiographical remembering (Angeli et al., 2025)^5^.

These observations raise questions about how processes related to target detection and mnemonic familiarity – separately or together – support recognition memory. Miller and colleagues recently examined response bias in neuroimaging studies of recognition memory and concluded that mnemonic familiarity signals interact with decision criteria to modulate neuronal responses to old items (Aminoff et al., 2015; Layher et al., 2023). Other studies showed that manipulating the old/new item ratio leaves posterior parietal old/new effects largely stable (see Table 2 of Herron et al., 2004), but produces probability-dependent differences in multiple prefrontal regions (Wagner et al., 1998; Herron et al., 2004), suggesting that the context of which retrieval processes happen matter. The present study sought to disentangle component processes that support recognition memory. To do so, we built on prior task designs by combining features of old-new recognition paradigms and visual oddball detection tasks. We first replicated the classical paradigms used in both research lineages, demonstrating that SAL/PMN is activated by old items in a traditional old-new recognition test and also by salient target events in a non-mnemonic oddball paradigm. We then directly contrasted these two processing dimensions (extending from Shannon & Buckner, 2004; see also Aminoff et al., 2015). Instead of presenting equal numbers of old and new words and requiring responses to both, our modigied paradigm introduced an imbalance. In one task condition, participants were presented mostly new words and only responded to the uncommon old words (Old Recognition). In another condition on another day, the same participants were presented mostly old words and only responded to the uncommon new words (New Recognition). In this manner, we separated mnemonic familiarity from target saliency.

## Methods and Materials

### Overview

The present study sought to (1) estimate the detailed functional network organization within intensively scanned individuals using resting-state functional connectivity MRI data and then (2) probe the response properties of the networks to determine the features of old / new recognition tasks that drive their response. Each of 6 participants underwent 4 separate MRI sessions (N=24 sessions). Thus, the study adopted an intensive within-subject design, as implemented previously in a growing series of studies from our laboratory (Braga et al., 2020; DiNicola, Braga, & Buckner, 2020; DiNicola, Sun, & Buckner, 2023; Du et al., 2024; Angeli et al., 2025), to target novel questions related to recognition memory. All data presented here are newly acquired and do not overlap previously reported studies.

### Participants

Six healthy right-handed adults aged 19 - 24 were recruited from the Boston area and paid for participation (M = 21.7, SD = 2.6, 2 male). Participants were English speakers (i.e., native or learned before age 7) and screened to exclude severe neurological or psychiatric illness. Participants provided informed consent using procedures approved by the Harvard University Institutional Review Board.

### MRI Data Acquisition and Preprocessing

MRI scans were conducted at the Harvard Center for Brain Science using a 3T Siemens MAGNETOM Prisma^fit^ MRI scanner with the vendor-supplied 32-channel phased-array head-neck coil (Siemens Heathineers AG, Erlangen, Germany). Head motion was controlled using foam and inflatable padding, and participants were instructed to remain still and alert. Visual stimuli were presented on a rear-projected display (F80-4K7 Laser Phosphor Projector; Barco, Kortrijk, Belgium) viewed via a mirror mounted to the head coil. Eye position and eyelid closure were video monitored (EyeLink 1000 Plus, Long-Range Mount; SR Research, Ottawa, Canada) and used to score alertness during each functional run.

Functional runs for both the resting-state and task-based acquisitions targeted blood oxygenation level-dependent (BOLD) contrast. A multiband gradient-echo EPI sequence (Feinberg et al., 2010; Setsompop et al., 2012; Xu et al., 2013) used 2.4-mm isotropic voxels, TR = 1,000 ms, TE = 33 ms, glip angle = 64°, matrix = 92 × 92, and 65 axial slices covering the cerebrum and cerebellum with a field of view of approximately 221 × 221 mm. The sequence was generously provided by the Center for Magnetic Resonance Research (CMRR) at the University of Minnesota. Dual-gradient-echo B0 fieldmaps with spatial resolution matched to the BOLD acquisitions were also acquired for distortion correction (TR = 295 ms, glip angle = 55°, TEs = 4.45 / 6.91 ms).

For anatomical registration and extraction of the cortical surface, a T1-weighted (T1w) multi-echo magnetization-prepared rapid acquisition gradient-echo (ME-MPRAGE; van der Kouwe et al., 2008) image was acquired using 0.8-mm isotropic resolution, matching the sequence protocol of the Human Connectome Project (HCP) (TR = 2,500 ms, TI = 1,000 ms, glip angle = 8°, TEs = 1.81/3.60/5.39/7.18 ms, matrix = 320 × 320 × 208, in-plane GRAPPA = 2). A spatially aligned T2-weighted (T2w) image was collected with a sampling perfection with application-optimized contrasts using different glip angle evolution (SPACE) sequence at 0.8-mm isotropic resolution (TR = 3,200 ms, TE = 564 ms, 208 slices, in-plane GRAPPA = 2). A backup (fast) T1w ME-MPRAGE image was also acquired using 1.2-mm isotropic resolution (TR = 2,200 ms, TI = 1,100 ms, glip angle = 7°, four echoes at TEs = 1.57/3.39/5.21/7.03 ms, matrix = 192 × 192 × 144, in-plane GRAPPA = 2).

Data were preprocessed using a custom pipeline designed to preserve fine-grained, individual-specific details by minimizing blurring and avoiding repeated resampling (“iProc”; Braga et al., 2019; DiNicola, Braga, & Buckner, 2020; Du et al., 2024). Brief method details are repeated here. The T1w 0.8-mm image was processed with FreeSurfer (v6.0, Fischl, 2012) *recon-all* to ensure accurate pial and white-matter boundaries and to establish a native-space target volume at 1.0-mm isotropic resolution. For each BOLD run, volumes were realigned to the run’s middle volume with FSL (v6.0.7, Jenkinson et al., 2012) *MCFLIRT* using 6 degrees of freedom, and the middle volume was unwarped using the dual-echo field map with FSL *FUGUE* to correct susceptibility-induced distortions. A participant-specific mean BOLD template was then constructed by up-sampling an unwarped low-motion middle volume to 1.2 mm as an interim target, registering the unwarped middle volumes from all runs to this target with affine transforms, and averaging the aligned images to yield a high-SNR mean BOLD image. The unwarped middle BOLD volume was registered to this mean BOLD template with affine transform using FSL *FLIRT* and 12 degrees of freedom, and the mean BOLD template was subsequently aligned to the participant’s T1-weighted image using boundary-based registration from FreeSurfer with 6 degrees of freedom.

All transforms were composed and applied in a single interpolation step to resample every BOLD volume directly into the 1.0-mm T1w native-space volume. This resampling strategy preserves spatial fidelity. For resting-state fixation data, nuisance regressors (e.g., 6 motion parameters, whole-brain signal, ventricular CSF signal, white-matter signal, and the first temporal derivative of each) were estimated from the native-space BOLD time series and removed with AFNI (Cox, 2012) *3dTproject*. The residuals were then band-pass filtered at 0.01–0.10 Hz using AFNI *3dBandpass*. For task data, only the whole-brain signal was regressed prior to model estimation, following prior work (DiNicola, Braga, & Buckner, 2020; DiNicola, Sun, & Buckner, 2023; Du et al., 2024, 2025; Angeli et al., 2025). After denoising, the BOLD data in native-volume space were sampled to the cortical surface and mapped to the *fsaverage6* template (40,962 vertices per hemisphere; Fischl, Sereno, & Dale, 1999) using FreeSurfer *vol2surf* projection. Surface data were smoothed with a 2-mm full-width at half-maximum Gaussian kernel using FreeSurfer *surf2surf*.

### Resting-State Fixation

Three to 4 resting-state fixation runs were collected per session, yielding a total of 14 to 16 runs per participant across sessions. Each run was 7 min 2 sec (422 frames, with the first 12 frames removed for T1 equilibration). Participants fixated a centered black plus sign displayed on an off-white background while remaining alert and still. The fixation runs were interleaved with active task runs (see *Active Task Paradigms* section), and the fixation data were used to estimate network organization through resting-state functional connectivity analysis. Importantly, the networks were estimated within each individual’s idiosyncratic anatomy prior to examination of task responses.

### Network Estimation Within Individuals

A multisession hierarchical Bayesian model (MS-HBM), initialized with a 15-network-parcellation prior, was used to estimate cerebral networks within individuals (see Du et al., 2024 for full procedure extended from Kong et al., 2019). For each participant, all available resting-state fixation runs were input to the model. Functional connectivity was computed at every surface vertex (40,962 vertices per hemisphere) on the *fsaverage6* cortical surface by correlating its BOLD timeseries with 1,175 uniformly distributed regions of interest across the *fsaverage* surface (Yeo et al., 2011), resulting in a 40,962 × 1,175 correlation matrix. For each vertex, the top 10% correlation coefficients were preserved to form a preliminary connectivity progile (Yeo et al., 2011; Du et al., 2024). The MS-HBM’s expectation maximization algorithm was then initialized with the 15-network group-level prior derived from the HCP S900 data release (Du et al., 2024), and individual connectivity profiles were entered into the model. Because our aim was to obtain the most accurate within-individual network estimates, only the model’s “training” stage was performed (as described in Du et al., 2024). All 6 participants were modeled jointly as one group. The 15 estimated networks were: Default Networks A (DN-A) and B (DN-B), Frontoparietal Networks A (FPN-A) and B (FPN-B), Somatomotor Networks A (SMOT-A), B (SMOT-B), Premotor-Posterior Parietal Rostral (PM-PPr), Dorsal Attention Networks A (dATN-A) and B (dATN-B), Visual Networks Central (VIS-C) and Peripheral (VIS-P), SAL/PMN, Cingulo-Opercular Network (CG-OP), Language Network (LANG), and an Auditory Network (AUD). See Supplemental Figure 1 for the full parcellation results across all participants.

The present analyses focused on two networks: SAL/PMN and CG-OP. SAL/PMN was the primary network of interest. SAL/PMN includes dACC, AI, and dorsolateral prefrontal cortex (dlPFC), which are key regions focused on in the literature referring to the network as the SAL network (Seeley, 2019; Seeley et al., 2007), as well as PCU, MCC, and IPL, which are the focus of the network’s description in the literature labeling it PMN (Gilmore, Nelson, & McDermott, 2015; McDermott et al., 2017; Gilmore, Nelson, et al., 2019). The separate names for the network – SAL and PMN – reflect two lineages that have converged on the same network, which is referred to here as SAL/PMN to reflect the historical origins (Du et al., 2024; see also Kwon et al., 2025; Ladwig et al., 2025). CG-OP was also included a target network as it contains adjacent regions in vAI that have been associated with responses to task-relevant, transient events (Dosenbach et al., 2006; Seeley, 2019; Du et al., 2024; see Dosenbach, Raichle, and Gordon, 2025 for discussion). More specifically, the Visual Oddball task used in this study (described below) robustly activates both the SAL/PMN and CG-OP networks (Du et al., 2024). Thus, CG-OP was included alongside SAL/PMN to capture network responses more fully to salient stimuli. Three additional higher-order networks were also included in post-hoc (exploratory) analyses that have been activated by effortful (controlled) processing (FPN-A), an adjacent right-lateralized network with uncertain functional properties (FPN-B; see Du et al., 2024), as well as network preferentially involved in episodic remembering / scene construction (DN-A).

### Model-Free Seed-Region-Based Verification of Network Estimates

To verify the SAL/PMN and CG-OP network estimates in each individual, model-free seed-region-based functional connectivity analyses were performed (e.g., Du et al., 2024). Pairwise Pearson correlations were computed between the fMRI time series of all surface vertices for each run. These correlations were Fisher z-transformed and then averaged across all runs to produce a stable within-subject connectivity matrix. This aggregated matrix was used to confirm the organization of the identified networks. Seed regions were manually placed in anterior and posterior zones within the MS-HBM-defined network boundaries. For SAL/PMN, seed regions were placed in the dlPFC (anterior) and the PCU (posterior); for CG-OP, seed regions were placed in the vAI (anterior) and lateral parietal cortex (posterior). These seed-region-based analyses independently confirmed the spatial patterns and internal consistency of the estimated SAL/PMN and CG-OP networks.

### Active Task Paradigms

During each MRI session, participants completed the same active task repeatedly. Across 4 independent MRI sessions, distinct tasks were performed allowing for robust estimates of the task effects within each person. The four tasks were designed to first replicate prior findings (Visual Oddball and Traditional Old-New Recognition) and then test functional response properties in an oppositional contrast, where recognition of old items was pitted directly against recognition of new items (Uncommon Old Recognition versus Uncommon New Recognition). The order of the 4 tasks was counterbalanced across participants to mitigate potential task order effects. Six runs were collected for each participant for each of the 4 tasks, for a total of 24 task runs collected per participant across the MRI sessions.

### Visual Oddball Task

The Visual Oddball task was designed to replicate the transient response to salient action-relevant events. During the task, participants were presented with a sequence of uppercase letters “O” and “K,” displayed in either black or red on an off-white background and were instructed to press a button using their right index finger when the red “K” appears, while withholding responses for all other letter-color combinations (Figure 1, right panel). This setup created a scenario where the red “K” served as the salient oddball target, and all other combinations acted as non-targets and distractors.

In each run, 10% of the trials featured the target red “Ks”, 10% were lure red “Os”, 40% were distractor black “Ks”, and 40% were distractor black “Os”. The contrast of interest was the target red “Ks” versus black distractor “Ks” and “Os”. The trial order within runs was randomized using Optseq (Dale, 1999) and tasks were implemented using PsychoPy (v2021.1.4). Each run for this and all task variations was 5 min 50 sec (350 frames, with the first 6 frames removed for T1 equilibration). The run structure included 6 sec of fixation overlapping with the initial stabilization frames, followed by a 20-sec fixation block, a 2-sec “Begin” cue, a continuous extended block of 150 trials (ITI = 2 sec; 500 ms letter followed by 1500 a black fixation dot), a 2-sec “End” cue, and then a final 20-sec fixation block. The trial duration was extended from 1 sec, as originally used in Du et al. (2024), to 2 sec to match the recognition task structure, minimizing task differences between the critical contrasts of interest in this study.

### Traditional Old-New Recognition

The Old-New Recognition task sought to obtain functional responses during a standard item recognition task where half of the items are old and half are new, and the participants respond to each item. Items were visually presented English words. Words were studied on the same day prior to the scanning session in a behavioral room adjacent to the scanner and then tested using old-new recognition during the MRI session. Word lists for this task (and the additional variations described below) all used the same corpus of matched lists. 108 unique 15-word lists of nouns were constructed, selected from the Corpus of Contemporary American English (Davies, 2010, December 2015 version). Lists were matched for word length (range = 3-7 letters, mean = 5.6 letters), frequency (range = 0.04-321 occurrences per million words; mean = 23.1 to 23.2 occurrences per million words across lists), and syllable count (range = 1-4 syllables; mean = 1.7 to 1.8 syllables across lists). The word lists were counterbalanced across participants and task versions such that words used as old in one recognition task for one participant would be used as new words in a different task version for another participant. This approach ensured that no specific word list was consistently associated with either familiarity or novelty, or any task variation.

During the pre-scan encoding, participants performed an incidental encoding task on 90 unique words that would become the old items during scanning (15 per run × 6 runs). The encoding task was repeated 4 times for each word, totaling 360 trials. Participants rated how much they liked the meaning of each word on a 5-point Likert scale. This encoding task, based on a pleasantness judgement, encouraged deep incidental encoding and has the feature that can be applied to any arbitrary word list (e.g., Squire et al., 1992; Buckner et al., 1995). By using multiple repetitions, the encoding phase intentionally encouraged a strong mnemonic signal. The encoding phase ended 20 to 40 min before scanning.

During the scanning phase, participants completed 6 runs of standard old / new recognition. Each run used 6 15-word lists with words from one list designated as old words and words from the remaining 5 lists designated as new words. Within a run, the 15 old words (studied during the pre-scan encoding phase) repeated 5 times, while the 75 new words (unfamiliar and not seen during encoding) were shown once (Figure 1, left panel), resulting in 150 trials. Thus, in any given presentation of the old word, the item had been seen at least 4 times previously and as many as 8 times. 50% of the trials were old words and 50% new words. Participants were instructed to press a button with their right index finger for old and right middle finger for new. The responses were extensively practiced prior to the first scan. The task contrast compared old words to new words. The run structure followed the same format as the oddball task in terms of run length, fixation lead in, and the cues to begin and end.

### Uncommon Old Word Recognition

The Uncommon Old Word Recognition task, which we refer to in short form as “Old Recognition”, combined salience (target) detection and recognition of familiar words. The structure and format of the task was similar to the traditional old / new recognition task but with two key differences. First, the participants responded only to old words, pressing with their right index finger (extending from Shannon & Buckner, 2004). Second, the old words were uncommon (10%) (Figure 2, left panel). Specifically, each run used 10 15-word lists, with a single list being old words and the remaining 9 lists being new words. In this way, the structure of this recognition task is also like the oddball task where the participant must respond to uncommon targets. Here though, the stimuli themselves were nominally the same across all trials (single words in the same color). The feature that differentiated the uncommon targets was their mnemonic history – the targets were targets because they were studied (familiar).

### Uncommon New Word Recognition

The Uncommon New Word Recognition task, which we refer to in short form as “New Recognition”, was designed as an oppositional contrast to the Old Recognition task. Here the targets were new words rather than old words. In this manner, when examining both the Old Recognition and New Recognition tasks, distinct response patterns can differentiate whether the familiarity or the saliency (target status) of the stimulus drives the response. The structure and format of the New Recognition task was identical to the Old Recognition task except the participants responded to uncommon new words (10%) (Figure 2, right panel). Each run used 2 15-word lists, with a single list being new words and the other list being old words that were repeated.

### Exclusion Criteria

Run exclusions followed previously used criteria (refer to DiNicola, Braga, & Buckner, 2020; Du et al., 2024; Xue et al., 2021) and were finalized prior to performing functional connectivity and task analysis. Four out of 239 acquired BOLD runs were excluded (1.7%). Three runs were excluded because maximum absolute motion was 1.90 or greater, and an additional run was excluded due to the head coil disconnecting. All runs were also examined for alertness through eye-tracking video monitoring and no runs were excluded. Overall, data quality was high with few exclusions.

### Replication of SAL/PMN and CG-OP Response in Oddball and Traditional Old / New Recognition Tasks

Analyses sought to replicate the response in SAL/PMN to familiar items in a traditional memory recognition task as well as the response to salient transients in a non-mnemonic oddball detection task. These are the two functional response properties that lead to the current debate about the role of SAL/PMN. Two questions were asked. First, given precise estimates of SAL/PMN within each individual participant, would a response be detected in one or both paradigms? Second, given that the literature on the SAL network has focused on vAI / dACC and the PMN network on PCU / MCC / IPL, would the topography of the responses include these hallmark regions in both kinds of task paradigm?

In addition to examining response properties in SAL/PMN, responses were also explored in CG-OP. CG-OP is adjacent to SAL/PMN in many zones of cortex, and SAL/PMN and CG-OP are challenging to anatomically and functionally distinguish as CG-OP also responds to task transients, especially when a motor action (response) is required (e.g., see Du et al., 2024 Figure 25 and Dosenbach, Raichle, and Gordon 2025). Thus, while the debate in the literature is primarily framed as focusing on SAL/PMN, many prior studies using group-averaged data and paradigms that do not dissociate the two partially adjacent networks are likely also reporting responses from CG-OP. For these reasons we examined CG-OP as an a priori target.

Task-related activity was modeled with a first-level general linear model (GLM) in SPM12 (https://www.gil.ion.ucl.ac.uk/spm/). For each participant, all available runs from each task were concatenated into a single time series. The design matrices used SPM’s canonical hemodynamic response function with a temporal derivative and a 100s high-pass filter, with additional run-level intercepts to accommodate baseline differences across runs. The GLM produced β-values and t-values for each surface vertex, which were subsequently converted to z-values using the *spm_t2z* function in SPM. For the Visual Oddball task, the GLM featured target red “Ks”, non-target black distractors, salient lure red “Os”, and cue onset/offset as regressors. Taking the difference between target red “Ks” and non-target black distractors z-statistical maps produced the Visual Oddball Effect. For Old-New Recognition, old words and new words were modeled as distinct regressors. The Old-New Recognition Effect was defined as old versus new words. Note that, as will be shown, performance accuracy was near ceiling and thus we chose to focus on the simplest form of analysis – old versus new items, without further subdivision into Hits versus CR, which we and others have done in situations where there is more variability in performance.

To explore the detailed topography of the response patterns and determine whether canonical SAL and PMN regions were activated including their multiple distributed regions, the individualized task response maps are presented along with the a priori defined boundaries of the networks. To quantify the effects in an unbiased manner, for each participant, mean z values were extracted from unthresholded contrast maps within individually defined boundaries of the SAL/PMN and CG-OP networks. These values were then averaged within each network for each task across both hemispheres, and group-level means, and standard errors were computed and plotted across the 6 participants. To assess whether activation within each network was significantly different from zero, one-sample t-tests were performed separately for SAL/PMN and CG-OP. In addition, the responses in FPN-A, FPN-B, and DN-A were also explored post-hoc without any specific a priori predictions regarding their response properties.

### Oppositional Test of SAL/PMN and CG-OP Response to Familiarity Versus Saliency

The Old Recognition and New Recognition tasks were designed to systematically oppose salience detection and familiarity-based recognition, affording a hypothesis-driven test of which component processing demand drives response in SAL/PMN. Specifically, a positive target effect within SAL/PMN for old items in the Old Recognition task would provide evidence consistent with both the salience and familiarity hypotheses. In contrast, a positive target effect within SAL/PMN for new items in the New Recognition task would support the salience hypothesis while refuting the mnemonic familiarity hypothesis, indicating that the SAL/PMN network is primarily driven by the detection of task-relevant target stimuli regardless of their mnemonic history. That is, if SAL/PMN responds to familiarity, it should not respond to target words that are new when they are contrast against a baseline of old, familiar words. Rather, a response to new words when they are targets would strongly refute a familiarity hypothesis and be consistent with a role of SAL/PMN in salience detection. CG-OP was also again explored given its anatomical adjacency and related functional properties.

### Group-Averaged Response Maps

While the primary design of the study focused on within-individual extraction of response estimates from individualized network estimates, we also created group-level contrasts. The study was not powered for a random effects group-level analysis, but the consistency of the effects and robustness of the maps are nonetheless informative as they provide a visualization of the group estimate that is similar to prior group-based neuroimaging studies of recognition memory. Task effects at the group level were created by mean averaging the individual z-statistical maps across participants for each task. This map should be considered a descriptive map of the average (gixed) effect (e.g., similar to Saadon-Grosman et al., 2022).

To provide group-level network references to aide visualization of the task-response maps, group consensus networks were also constructed by assigning each vertex the network label that appeared most often in the individually parcellated MS-HBM maps across six participants (similar to Supplemental Figure 18 in Du et al., 2024). Isolated holes were gilled and incontiguous clusters that contained 15 or fewer voxels were removed using Connectome Workbench (v1.3.2, Marcus et al., 2011). The surviving clusters were then combined to produce the final group-level consensus estimates of the networks. These group-level network estimates provide a rough estimate of each network’s position in the group given between-subject anatomical variability. Small, variable regions, such as in IPL, will have minimal consensus. Thus, these group network references are consensus estimates of most consistent locations of regions. They are not valid estimates of the extent or detailed borders of regions as would appear in any given individual.

## Results

### Old-New Recognition Reveals Asymmetric Response Patterns

Recognition accuracy in the Old-New Recognition task was near ceiling (Figure 3A). There were extremely few Misses or False Alarms (FA) consistent with a strong familiarity signal. The mean Hit-FA rate across participants was 0.94 with a range of 0.90 to 0.99. Mean response times revealed that CR were slower than Hits by 55 msec (Figure 3B). Detailed examination of the response time distributions for each participant supported this asymmetrical effect (Figure 4). While the response time distributions were highly overlapping, most participants exhibited clear asymmetry, with response time distributions to new words visibly shifted rightward (i.e., slower) relative to old words. A Kolmogorov-Smirnov (K-S) test was performed on the response time distributions revealing significant effects in 5 of the 6 participants (all *p* < .01; S4 did not show a significant effect, *p* = .71).

**Figure 3.**
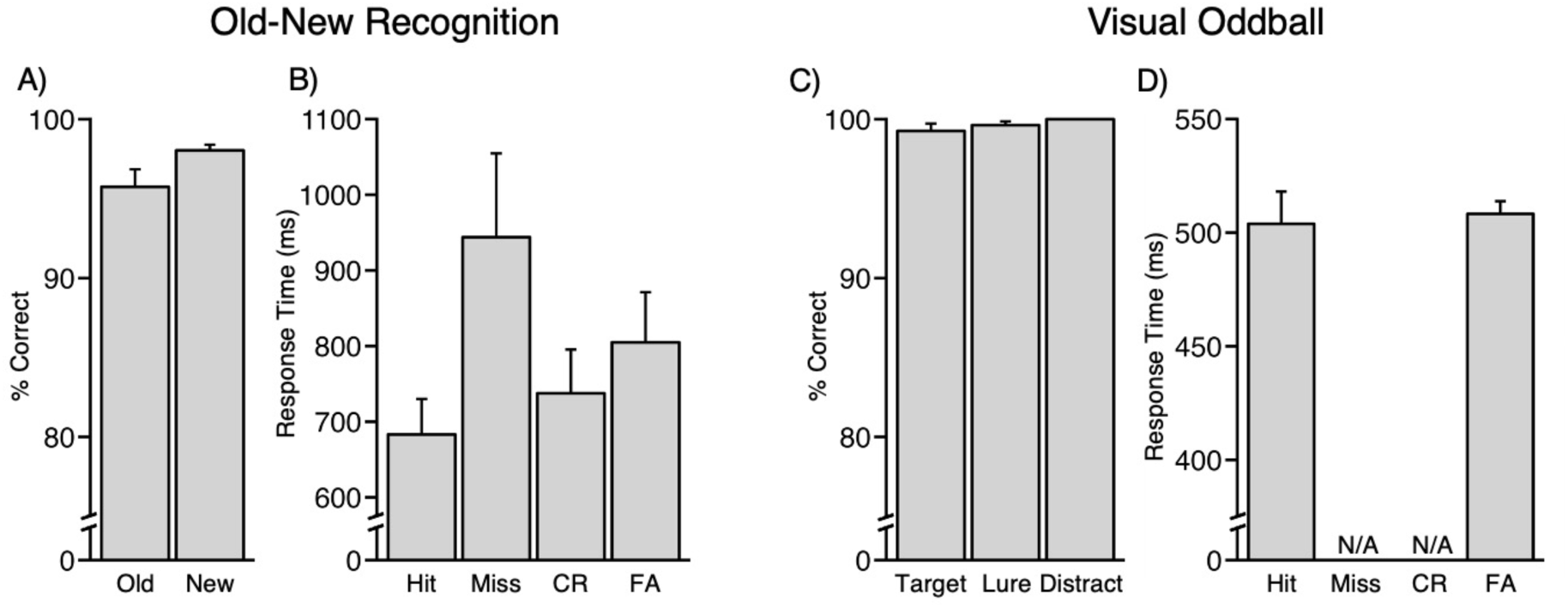
Behavioral results for the Old-New Recognition and Visual Oddball tasks. The mean performance and response times are shown for the traditional Old-New Recognition and Visual Oddball tasks. (A) Recognition performance was extremely high during Old-New Recognition. (B) Response times were fast during Old-New Recognition with the typical pattern of slower responses for Correct Rejections (CR) as contrast to Hits. False alarms (FA) and Miss responses were rare (see text) but their values are also plotted. (C) Target detection performance was also extremely high in the Visual Oddball task. (D) Response times for target detection were also fast. Error bars represent the standard error of the mean across participants.

**Figure 4.**
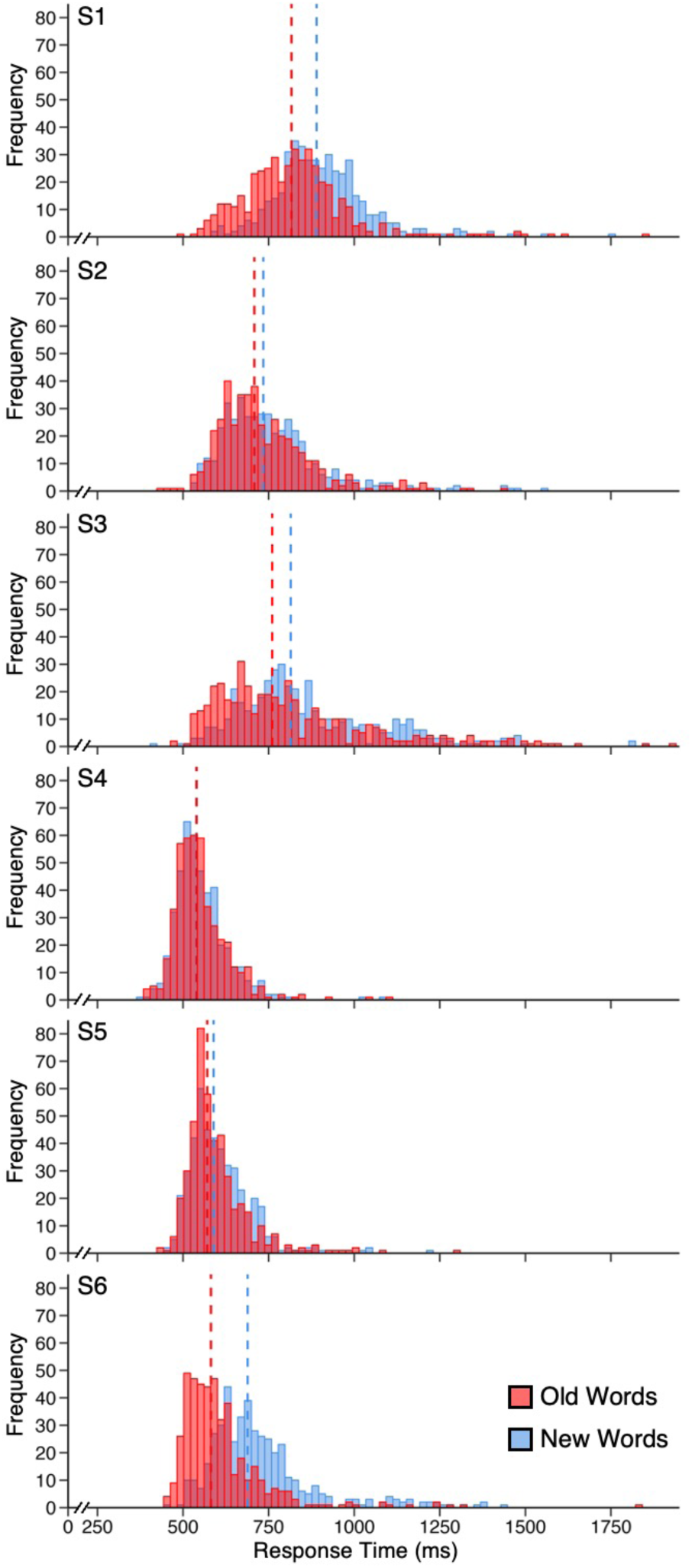
Response time distributions reveal old responses are faster than new responses in the Old-New Recognition task. Histograms show trial-level response times for old (red) and new (blue) words for each of the 6 participants. Each distribution includes responses aggregated across all runs. The dashed lines show the median responses. Statistical tests revealed the effect was significant within each participant except S4 (see text). Participants S1, S3, and S6 show the clear response time asymmetry, with the responses to new words visibly shifted to the right.

Target detection performance in the Visual Oddball task was also near ceiling (Figure 3C). All participants correctly withheld responses to the distractor black letters (i.e., no button presses were recorded). Response times were recorded only for trials that required a button press (Figure 3D). No response times were available for missed target red “Ks” (Miss) or correctly rejected lure red “Os” and distractor black letters (CR), as these did not require a response. Only two participants produced a single FA to either lure red “Os” or distractor black letters.

### SAL/PMN and CG-OP are Present and Similarly Organized in Each Participant

Both the SAL/PMN and CG-OP networks were estimated within each individual participant replicating anatomical features that are now established in the literature. Figures 5 and 6 display the network estimates from S1 and S6. Network estimates from all other participants are available in the Supplementary Materials.

**Figure 5.**
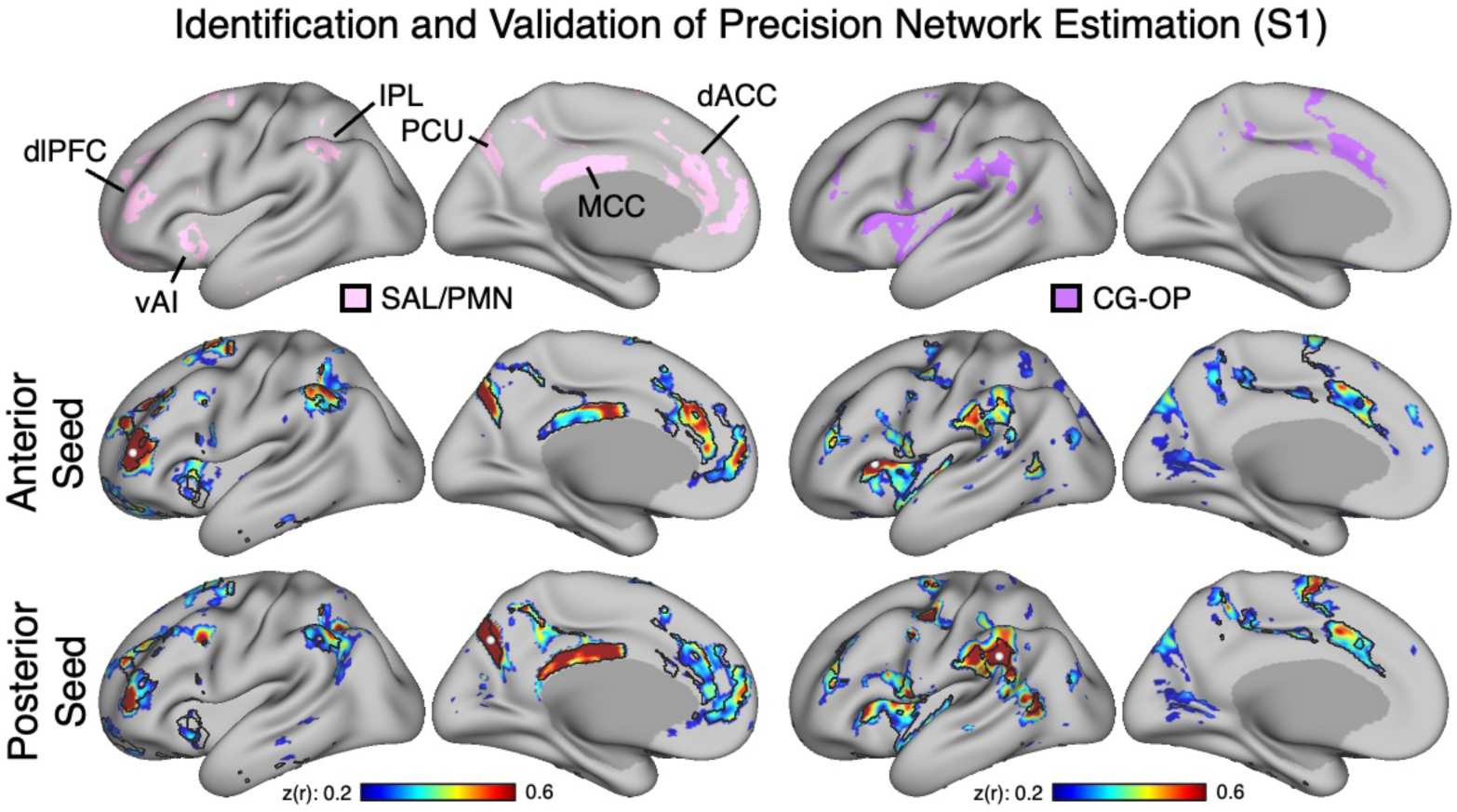
Target networks identified in S1. The SAL/PMN and CG-OP networks were automatically identified using a multisession hierarchical Bayesian model (MS-HBM) in participant S1 and then validated in the raw correlation data using seed-based analyses. (Top Row) The SAL/PMN (pink, left) and CG-OP (purple, right) networks are displayed in lateral and medial views of the inflated left hemisphere. (Middle Row) Correlation maps using anterior seed regions (white circles) are displayed for each network. (Bottom Row) Correlation maps using posterior seed regions (white circles) are displayed for each network. The black outlines show the boundaries of the automatically define network estimates. Correlation maps are plotted on a z(r) scale, with the color bar at the bottom. Note how the raw correlation patterns are spatially selective for the estimated networks suggesting the automated MS-HBM-derived network estimates capture the spatial correlation structure of the data. dlPFC, dorsolateral prefrontal cortex; vAI, ventral anterior insula; IPL, inferior parietal lobule; PCU, precuneus; MCC, mid-cingulate cortex; dACC, dorsal anterior cingulate cortex.

**Figure 6.**
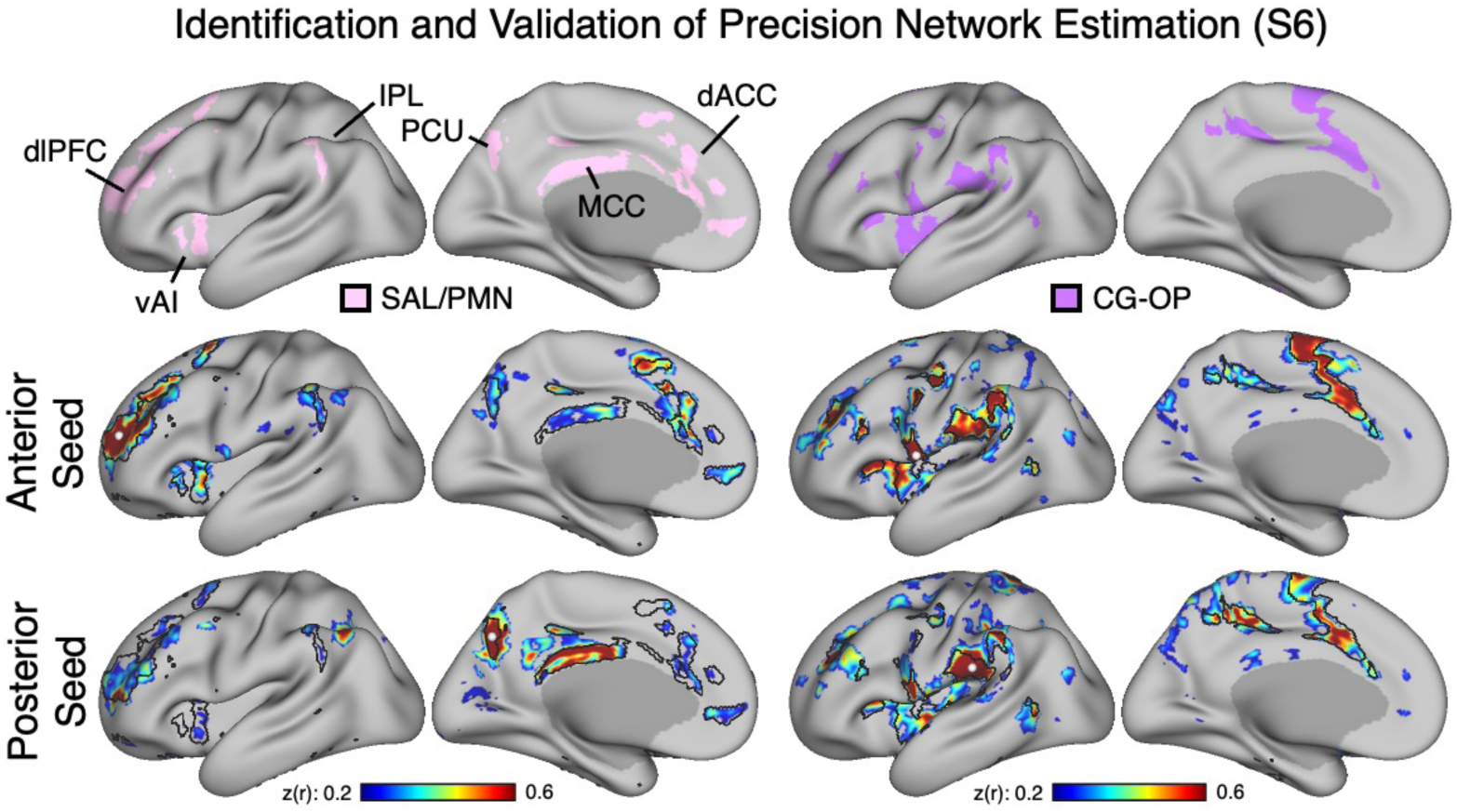
Target networks identified in S6. The SAL/PMN and CG-OP networks were automatically identified using a multisession hierarchical Bayesian model (MS-HBM) in participant S6 and then validated in the raw correlation data using seed-based analyses. (Top Row) The SAL/PMN (pink, left) and CG-OP (purple, right) networks are displayed in lateral and medial views of the inflated left hemisphere. (Middle Row) Correlation maps using anterior seed regions (white circles) are displayed for each network. (Bottom Row) Correlation maps using posterior seed regions (white circles) are displayed for each network. The black outlines show the boundaries of the automatically define network estimates. Correlation maps are plotted on a z(r) scale, with the color bar at the bottom. Note how the raw correlation patterns are again spatially selective for the estimated networks suggesting the automated MS-HBM-derived network estimates capture the spatial correlation structure of the data. dlPFC, dorsolateral prefrontal cortex; vAI, ventral anterior insula; IPL, inferior parietal lobule; PCU, precuneus; MCC, mid-cingulate cortex; dACC, dorsal anterior cingulate cortex.

Specifically, SAL/PMN included vAI and dACC as well as regions along the posterior midline (PCU and MCC) and IPL, thus revealing the canonical regions that have been the focus of both the literatures on the SAL and PMN networks, consistent with the hypothesis that there is only a single network. This distributed pattern could be fully reproduced with either anterior or posterior seed regions indicating that the network estimates were robust and not obligated by the use of spatial priors in the MS-HBM. Rather, the pattern could be reliably generated in every participant from the raw correlation data without prior assumptions about the network’s organization.

CG-OP was also reliably identified across all participants. While sharing a general organization in common with SAL/PMN and multiple side-by-side adjacencies, CG-OP differed in that its regions surrounded somatomotor cortex along the midline and laterally extending into the Sylvian fissure (see glat maps in Du et al. 2024). Seed region correlation patterns again confirmed the MS-HBM estimates with the additional observation that the correlations also included regions in occipital cortex as previously noted (see Du et al., 2024 Figure 22) as well as extensions into the precentral gyrus that form discontinuities in the body map (recently labeled inter-effector regions; Gordon et al., 2023).

These initial results confirm that the present precision mapping methods were able to robustly identify the core components of the SAL/PMN and CG-OP networks within each individual and further that the identification of the networks is not dependent on assumptions of the MS-HBM model. Having estimated the networks within individuals, we next turned to exploring the component processes that elicit responses within them.

### SAL/PMN Responds Modestly to Old Items During Recognition Memory and Robustly to Salient Targets in the Non-Mnemonic Oddball Task

#### Old-New Recognition

SAL/PMN displayed a positive response to old words compared to new words in the traditional Old-New Recognition task (Old-New Recognition Effect). Quantitatively, the Old-New Recognition Effect was present in all 6 participants to varying degrees (Figure 7A). Spatially the response showed a broad distribution that overlapped the core SAL/PMN regions in most participants including the posterior midline regions PCU and MCC (Figure 7B). The posterior response in the IPL was considerably more extensive than might be expected from a response in SAL/PMN alone. While the sample size was small, the positive response within SAL/PMN was significant at the group level in addition to being individually present within each participant [Figure 8B; *t*(5) = 3.54, *p* = .02].

**Figure 7.**
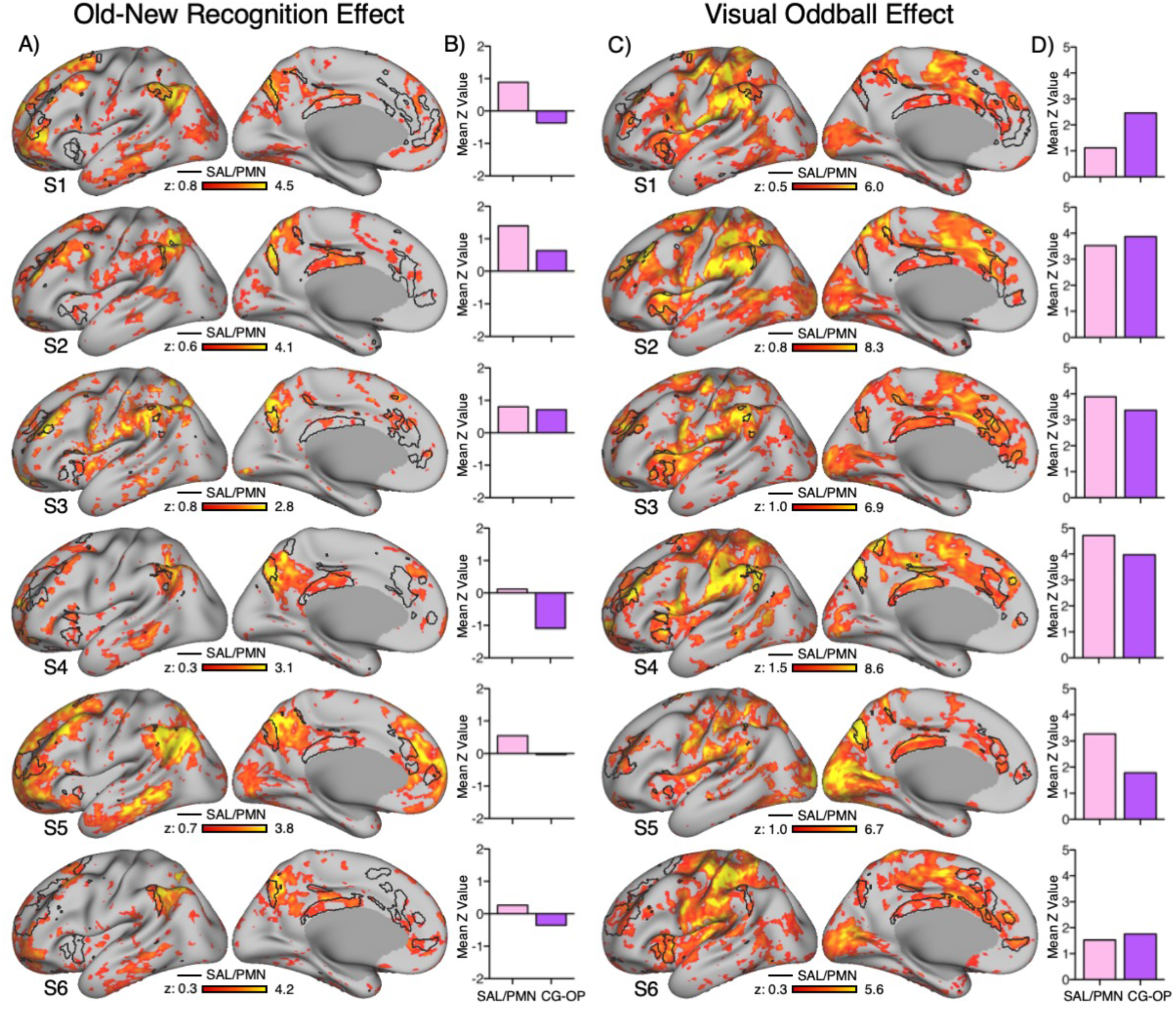
The Salience/Parietal Memory Network (SAL/PMN) responds during traditional Old-New Recognition and Visual Oddball tasks. Each row shows the task effects within an individual participant. (A) Old-New Recognition Effect maps are plotted on a z scale, with the color bar at the bottom. Each participant’s SAL/PMN network is outlined in black. (B) The Old-New Recognition Effect estimates within the automatically defined SAL/PMN and CG-OP networks are plotted (pink, SAL/PMN; CG-OP, purple). (C) Old-New Recognition Effect maps are plotted on a z scale, with the color bar at the bottom. Each participant’s SAL/PMN network is outlined in black. (C) Visual Oddball Effect maps are plotted on a z scale, with the color bar at the bottom. Each participant’s SAL/PMN network is again outlined in black. (D) The Visual Oddball Effect estimates within the automatically defined SAL/PMN and CG-OP networks are plotted (pink, SAL/PMN; CG-OP, purple). Note that a response pattern is observed for both task contrasts that overlaps the SAL/PMN network. The response is not selective for the network but includes, in both cases, the precuneus and mid-cingulate cortex.

**Figure 8.**
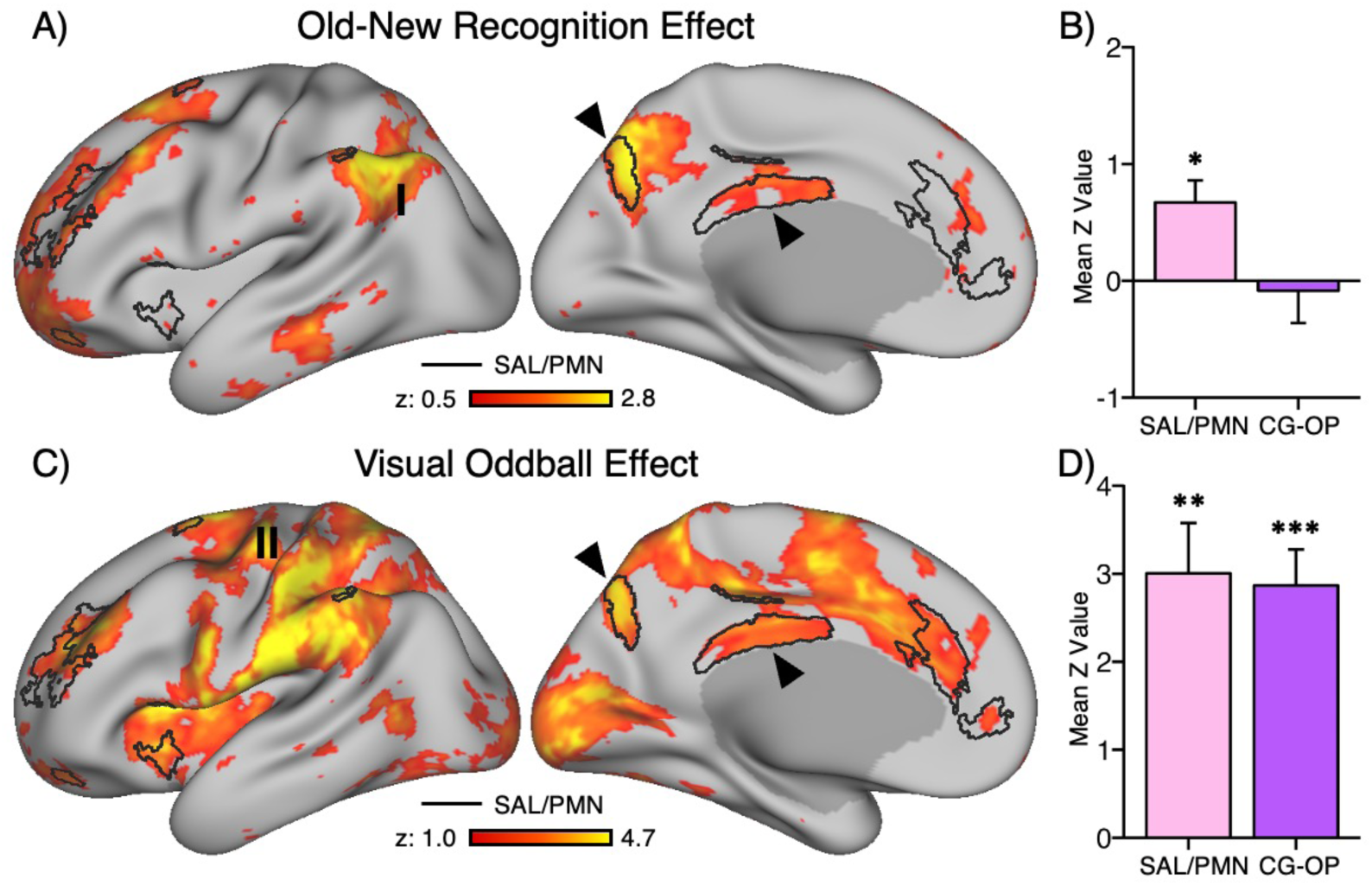
Group-level analyses confirm the Salience/Parietal Memory Network (SAL/PMN) responds during traditional Old-New Recognition and Visual Oddball tasks. Each row shows the task effect for the average group of participants. (A) The Old-New Recognition Effect map is plotted on a z scale, with the color bar at the bottom. These are fixed effect maps showing the mean Z across participants (hence the relatively low Z values). Thus, the maps are descriptive (not hypothesis testing) and reveal the average spatial topography of the response. The group-estimate SAL/PMN network is outlined in black. The arrows show the precuneus and mid-cingulate cortex regions. (B) The group-level means of the Old-New Recognition Effect estimates are plotted for the SAL/PMN and CG-OP networks (pink, SAL/PMN; CG-OP, purple). (C) The Visual Oddball Effect map is plotted on a z scale, with the color bar at the bottom. The group-estimate SAL/PMN network is outlined in black. The arrows show the precuneus and mid-cingulate cortex regions. (D) The group-level means of the Visual Oddball Effect estimates are plotted for the SAL/PMN and CG-OP networks (pink, SAL/PMN; CG-OP, purple). Note that the group effects reveal clear response in the SAL/PMN including the precuneus and mid-cingulate cortex in both contrasts. There is also a clear response in motor cortex (II) that likely reflects the right-handed keypress, as well as an unexpectedly extensive response in the inferior parietal lobule (I) that will be the focus of later analyses. Error bars represent the standard error of the mean across participants. * p < .05; ** p < .01; *** p < .001

By contrast, CG-OP was not (or minimally) activated by the Old-New Recognition Effect. The core regions along the precentral gyrus and surrounding the somatomotor cortex were absent. Four of the 6 participants showed a negative response at the network level and the (underpowered) group-level effect was not significant [*t*(5) = -0.3, *p* = .77]. Thus, while we cannot assert the null, CG-OP was at best minimally active. This minimal response will stand in stark contrast to the task contrasts below that have a differential motor response and strongly activate CG-OP.

Group-level maps of the Old-New Recognition Effect reinforced all of these observations and were similar to maps reported in the literature showing old / new recognition effects (Figure 8A). Particularly relevant to the recent literature on the PMN network, the response along the posterior midline with its canonical features was replicated as indicated by black arrows (e.g., see the clear response in PCU and MCC). An extended response in IPL was again observed at the group level (label I in Figure 8A). Note that the individual-level network regions for the SAL/PMN network are variable in IPL, leading to a small consensus region. That notwithstanding, the Old-New Recognition effect yielded a spatially extensive response in lateral parietal cortex, an anatomical feature we will return to later.

#### Visual Oddball

SAL/PMN and CG-OP both showed strong positive responses to targets in the Visual Oddball task relative to non-target distractors (Visual Oddball Effect; Figure 7D). The Visual Oddball Effect was present for both networks in all 6 participants. The response spatially overlapped the core SAL/PMN regions in every individual including the posterior midline regions PCU and MCC that historically define the PMN (Figure 7C). The effect was significant at the group level for both SAL/PMN [*t*(5) = 5.26, *p* < .005] and CG-OP [*t*(5) = 6.98, *p* < .001] networks (Figure 8D).

The Visual Oddball Effect in the group included spatially restricted positive responses in PCU and MCC, as illustrated by arrowheads in Figure 8C, in addition to responses within the CG-OP network including regions surrounding somatomotor cortex. There was also a clear response in left motor cortex near the hand representation consistent with the right-handed keypress response to targets (reference trials were absent a motor response) (labeled II in Figure 8C). Thus SAL/PMN and CG-OP, including their multiple canonical regions, were both robustly activated during target detection in addition to a broader array of regions consistent with the sensory and motor demands of the task.

### Behavioral Results for Old Recognition and New Recognition

Mean recognition accuracy in the Old Recognition task was 85.7% for old words and 99.4% for new words (Figure 9A). The mean Hit-FA rate across participants was 0.85 with a range of 0.72 to 0.93. Mean response times revealed that FA were slower than Hits by 205 msec (Figure 9B). Mean recognition accuracy in the New Recognition task was 91.3% for new words and 99.3% for old words and (Figure 9C). The mean Hit-FA rate across participants was 0.91 with a range of 0.80 to 0.98. Note that here, unlike typical old / new recognition tasks, Hits are correct responses to New, not old, items. Mean response times revealed that FA were again slower than Hits by 90 msec (Figure 9D). These behavioral patterns indicate that extremely high recognition performance was maintained, suggesting a strong mnemonic familiarity signal, while the salient target for detection was shifted from Old to New items between the two task variations.

**Figure 9.**
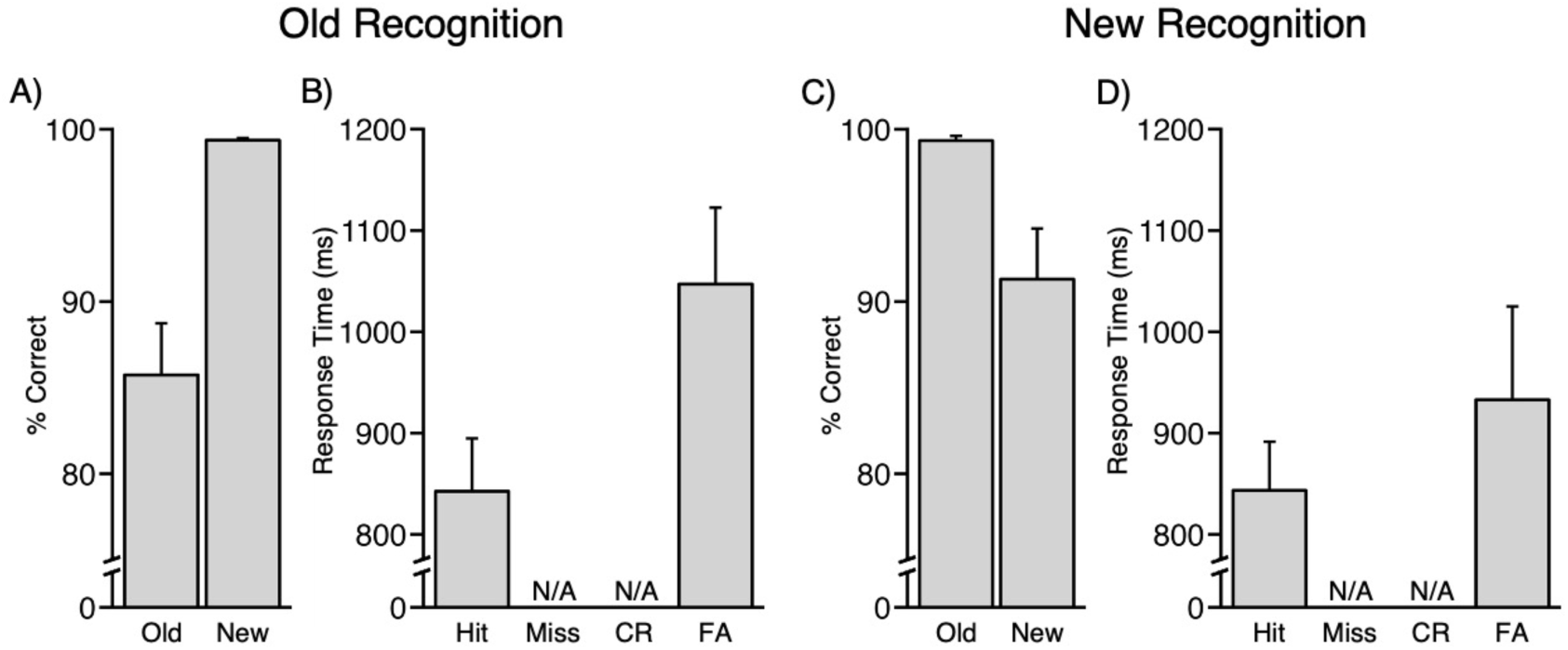
Behavioral results for the Old Recognition and New Recognition tasks. The mean performance and response times are shown for the Old Recognition and New Recognition tasks. (A) Recognition performance was extremely high during Old Recognition. (B) Response times were fast. Note that there was no required response for new items and thus no misses or correct rejections (CR). (C) Recognition performance was extremely high during New Recognition. (B) Response times were again fast. Note that here hits are correct responses to new, not old, items. Error bars represent the standard error of the mean across participants. CR, Correct Rejections; FA, False Alarms.

### SAL/PMN Responds to Old Items When They are Uncommon Targets

SAL/PMN displayed a positive response to old word targets compared to the new word distractors (Old Recognition Effect) in every individual as was a response in CG-OP (Figure 10B). The spatial extent of the Old Recognition Effect robustly included the posterior midline regions PCU and MCC (S5 did not show a clear response in MCC) (Figure 10A) as well as an extended set of regions similar to the pattern observed in the Visual Oddball Effect (Figure 7B). The present contrast contained an even more extensive pattern of response, possibly because the present task condition combined two types of effect: the recognition of familiar items and the detection and response to salient stimuli. The response in SAL/PMN [*t*(5) = 9.14, *p* < .001] and CG-OP [*t*(5) = 6.79, *p* <.001] showed significant positive responses to old items at the group level (Figure 11B).

**Figure 10.**
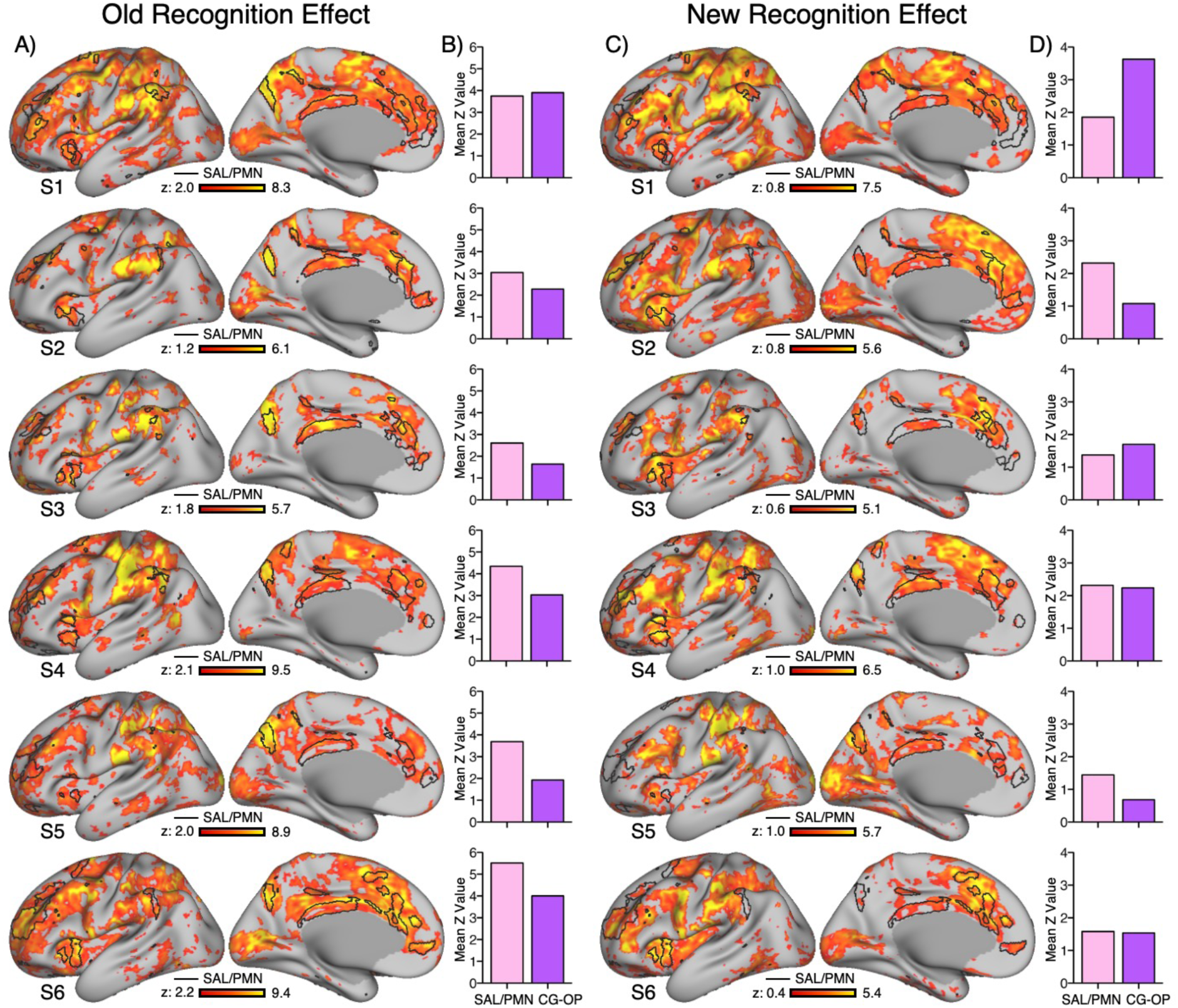
The Salience/Parietal Memory Network (SAL/PMN) responds to salient targets even when they are new items. Each row shows the Old Recognition Effect and New Recognition Effect within an individual participant. (A) Old Recognition Effect maps are plotted on a z scale, with the color bar at the bottom. Each participant’s SAL/PMN network is outlined in black. (B) The Old Recognition Effect estimates within the automatically defined SAL/PMN and CG-OP networks are plotted (pink, SAL/PMN; CG-OP, purple). (C) New Recognition Effect maps are plotted on a z scale, with the color bar at the bottom. Each participant’s SAL/PMN network is outlined in black. (D) The New Recognition Effect estimates within the automatically defined SAL/PMN and CG-OP networks are plotted (pink, SAL/PMN; CG-OP, purple). Note that a response pattern is observed for both task contrasts that overlaps the SAL/PMN network. Critically, the New Recognition Effect yields a response in canonical precuneus and mid-cingulate cortex regions in each participant, with some participants revealing robust spatially localized responses.

**Figure 11.**
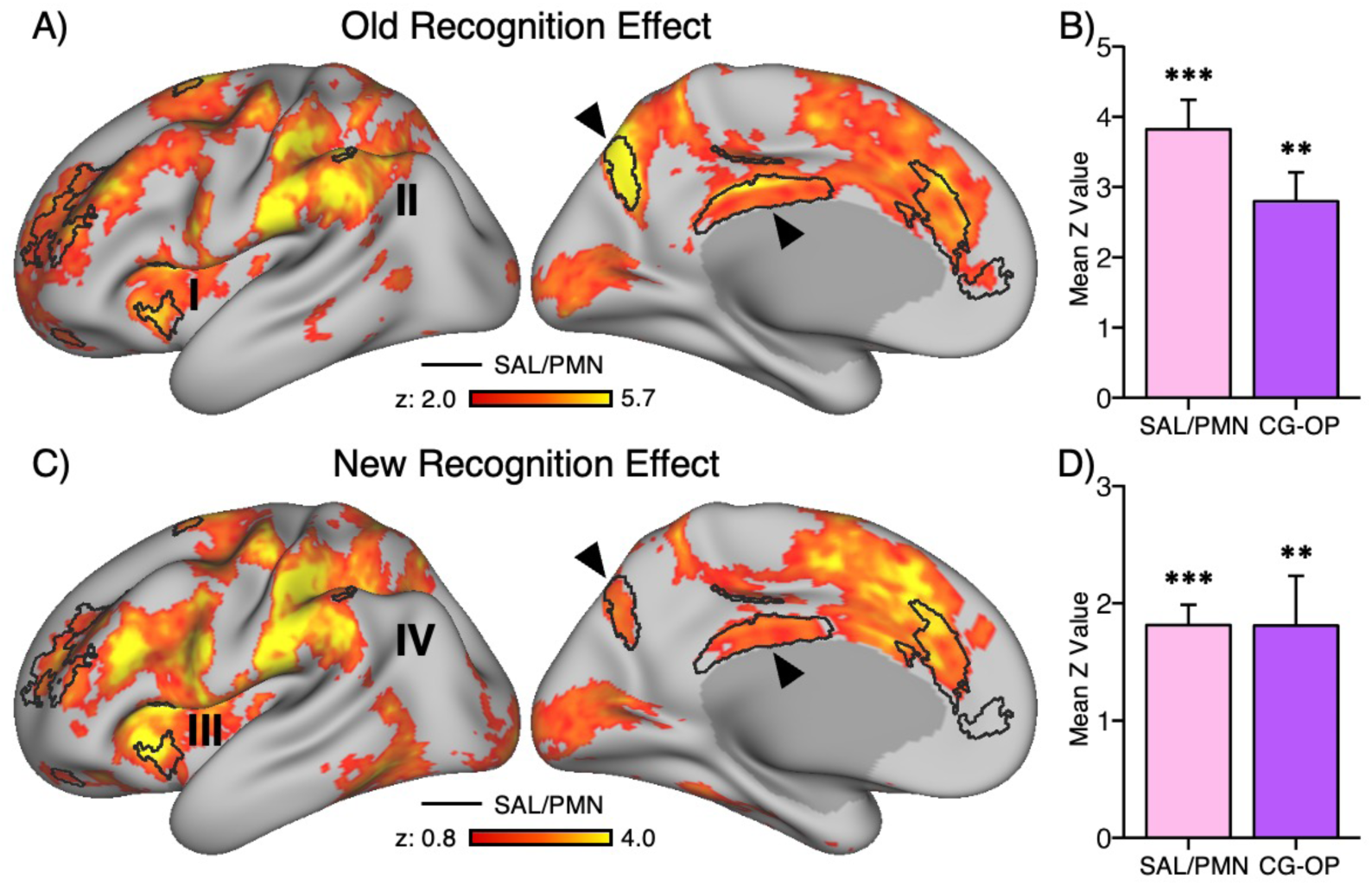
Group-level analyses confirm the Salience/Parietal Memory Network (SAL/PMN) responds to salient new targets. Each row shows the task effect for the average group of participants. (A) The Old Recognition Effect map is plotted on a z scale, with the color bar at the bottom. These are fixed effect maps showing the mean Z across participants. Thus, the maps are descriptive (not hypothesis testing) and reveal the average spatial topography of the response. The group-estimate SAL/PMN network is outlined in black. The arrows show the precuneus and mid-cingulate cortex regions. Note also the clear response in the anterior insula (I) and the extensive response in the inferior parietal lobule (II). (B) The group-level means of the Old Recognition Effect estimates are plotted for the SAL/PMN and CG-OP networks (pink, SAL/PMN; CG-OP, purple). (C) The New Recognition Effect map is plotted on a z scale, with the color bar at the bottom. The group-estimate SAL/PMN network is outlined in black. Note again the clear response in the anterior insula (III) but the more restricted response in the inferior parietal lobule (IV) as compared to the Old Recognition Effect map. (D) The group-level means of the New Recognition Effect estimates are plotted for the SAL/PMN and CG-OP networks (pink, SAL/PMN; CG-OP, purple). Error bars represent the standard error of the mean across participants. * p < .05; ** p < .01; *** p < .001.

Group-level maps of the Old Recognition Effect again reinforced the observations made in the individual participants (Figure 11A). Widespread response throughout the entire SAL/PMN network was observed covering all regions within the network boundary.

### SAL/PMN Also Responds to New Items When They are Uncommon Targets

To directly test whether SAL/PMN responses are driven by familiarity or salience, the New Recognition task was examined. In this task, uncommon salient new words were treated as the targets rather than familiar old words. Contrasting the new words against the old word distractors (New Recognition Effect) provides the critical oppositional contrast.

The spatial extent of the New Recognition Effect clearly included SAL/PMN. PCU and MCC were evident in nearly every individual, with the PCU and MCC regions of SAL/PMN showing a robust, spatially circumscribed response in several participants (e.g., S4). The response was not restricted to SAL/PMN, but SAL/PMN was clearly responding to new items when they were targets against a backdrop of familiar items, a pattern that cannot be accommodated within a simple item repetition or familiarity hypothesis. CG-OP was also clearly active, paralleling the observation across all contrasts in the present study where the target condition included an active motor response, and the reference condition did not.

Group-level analyses revealed a significant positive response in both SAL/PMN [t(5) = 10.5, p < .001] and CG-OP [t(5) = 4.27, p < .01] networks (Figure 11D). The group-level map of the New Recognition Effect showed widespread responses across SAL/PMN, closely resembling patterns observed in the Old-New Recognition Effect (Figure 8A) and Visual Oddball Effect (Figure 8C), apart from reduced involvement in IPL and also regions extending across lateral PFC along the inferior frontal gyrus (Figure 11C marked IV). We explore these exceptions in post-hoc exploratory analyses in the next section.

### Exploratory Analysis of the Familiarity Effect

The hypothesis-directed explorations above separated processes linked to salience (target) detection and familiarity. Both the individual-level and group-level analyses supported that SAL/PMN responds when targets are salient, regardless of their mnemonic familiarity. Thus, the current results argue against a specific contribution of SAL/PMN to familiarity-based recognition. However, the results noted to this point offer little insight into what networks might preferentially support recognition memory. In a final post-hoc set of analyses, that should be considered idea raising, we revisited all task contrasts to ask which network (or networks) might preferentially support mnemonic familiarity.

A familiarity-based response would be expected to be present when old items were contrasted to new items, regardless of target status (here those contrasts would include the Old-New Recognition Effect and the Old Recognition Effect). Figure 12 shows all contrasts for participants S1 and S6 with the right hemisphere displayed to reveal an intriguing, unexpected effect: FPN-B, a right-lateralized network, was active in the Old-New Recognition Effect and the Old Recognition Effect contrasts but not the New Recognition Effect contrasts. The Supplementary Materials show similarly plotted results for all participants. Figure 13 quantigies this effect contrasting the behavior of FPN-B with FPN-A (a network that responds consistently during effortful tasks regardless of domain) and DN-A (a hippocampal-linked network that responds during episodic remembering especially when the process involves scene construction). FPN-B reveals a robust response consistent with a familiarity (oldness) signal (Figure 13B, F).

**Figure 12.**
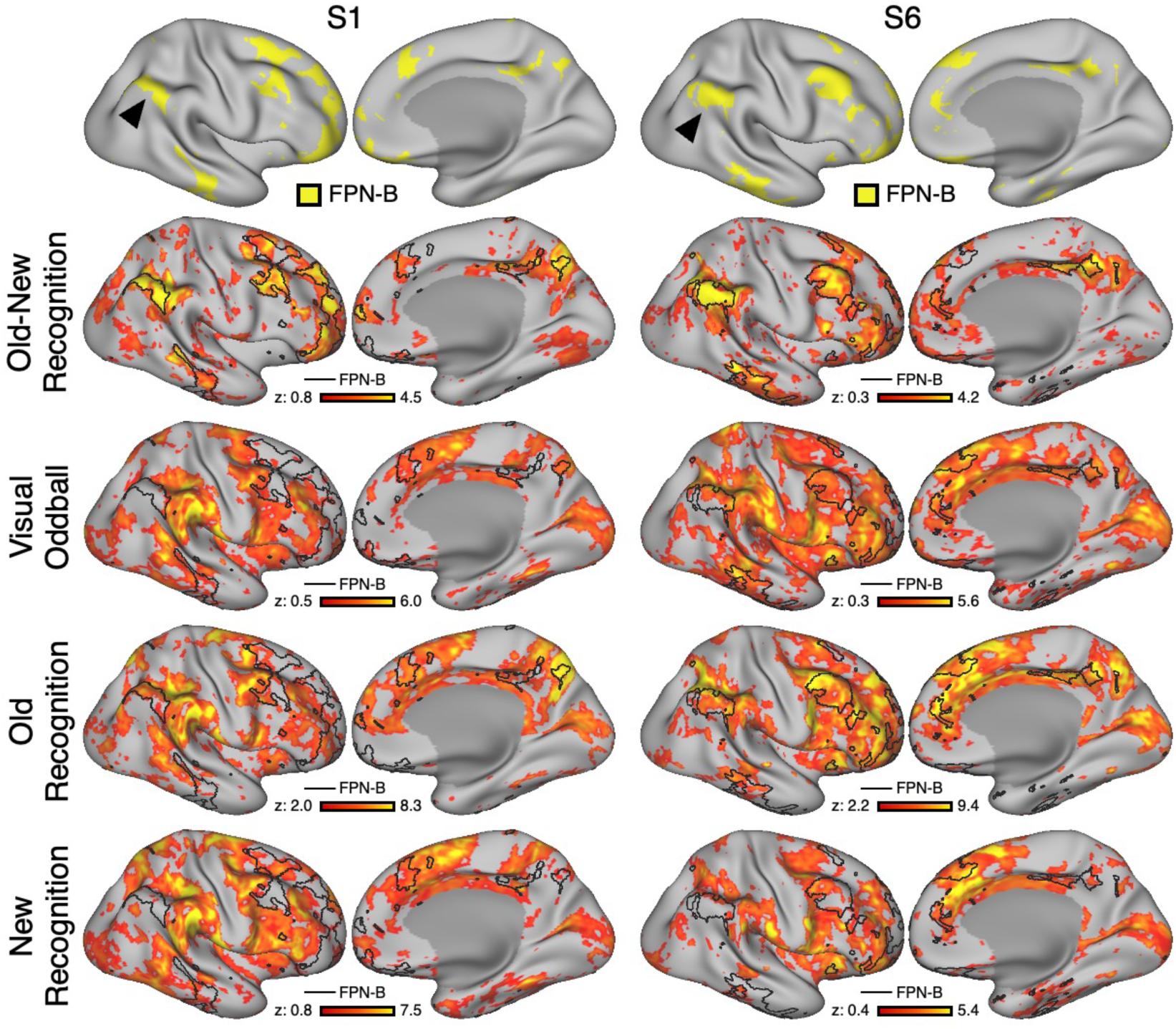
Exploratory analyses suggest Frontoparietal Network B (FPN-B) responds to familiarity. (Top Row) FPN-B (yellow) is displayed in lateral and medial views of the inflated right hemisphere for two representative participants (S1, S6). The right hemisphere is displayed because FPN-B is strongly right lateralized. Note the FPN-B consistently includes a region in caudal inferior parietal lobule in all participants. (Rows Two to Five) Old-New Recognition, Visual Oddball, Old Recognition, and New Recognition Effect maps are displayed for both representative participants. Effect maps are plotted on a z scale, with the color bar at the bottom. Each participant’s FPN-B is outlined in black. Note that the Old-New Recognition Effect response falls within the distributed regions of FPN-B prominently including the FPN-B portion of the inferior parietal lobule. The effect is also present in the Old Recognition Effect map but is absent in the Visual Oddball and New Recognition Effect maps. The Supplemental Materials show maps for both hemispheres for all participants.

**Figure 13.**
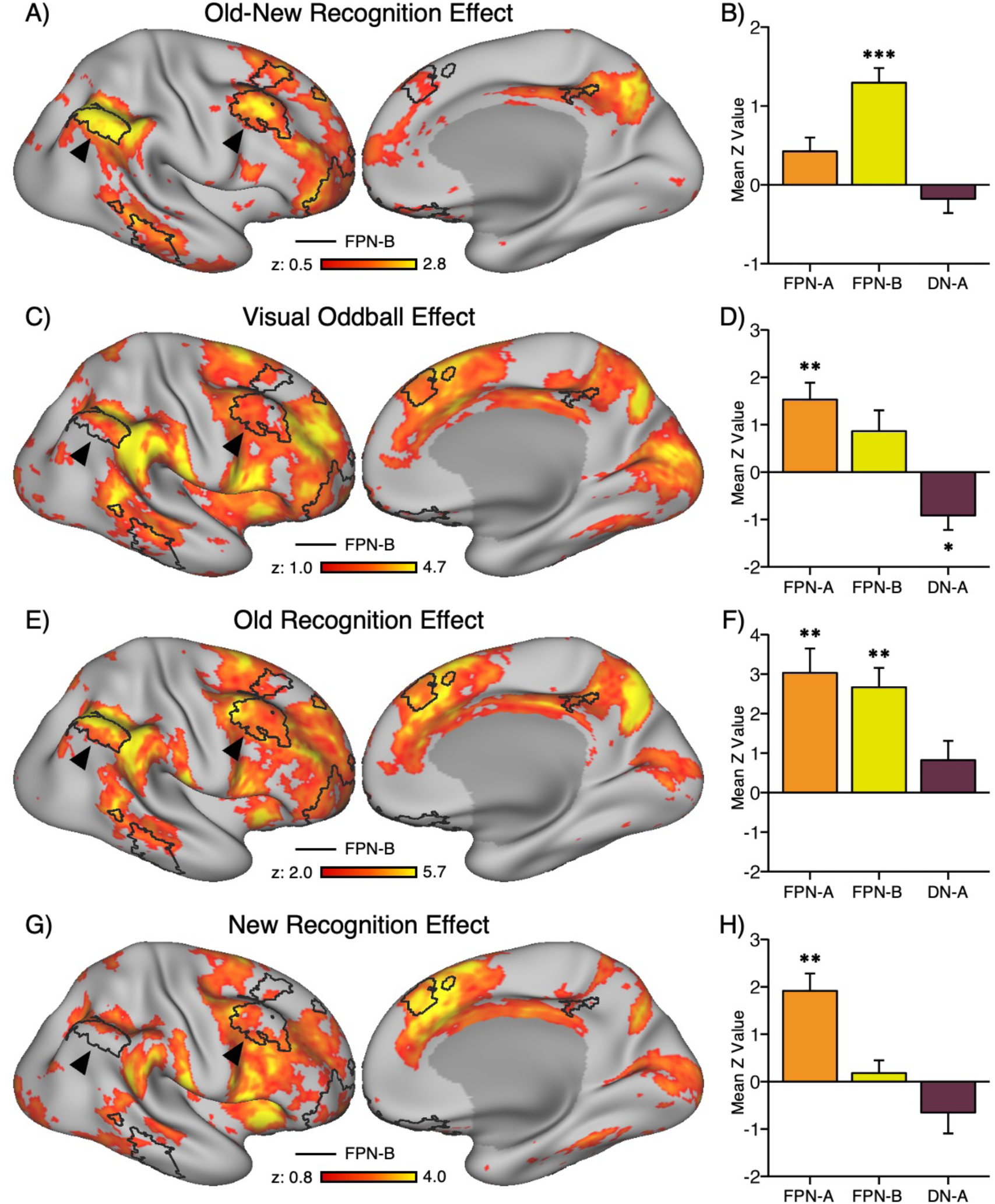
Group-level analyses confirm the Frontoparietal Network B (FPN-B) responds to familiarity. Each row shows the task effect for the average group of participants. (A) The Old-New Recognition Effect map is plotted on a z scale, with the color bar at the bottom. The group-estimate FPN-B is outlined in black. The arrows show the FPN-B regions of the caudal inferior parietal lobule and dorsolateral prefrontal cortex. Note the clear response in the inferior parietal lobule. (B) The group-level means of the Old-New Recognition Effect estimates are plotted for the Frontoparietal Network A (orange, FPN-A), FPN-B (yellow), and Default Network A (brown). The response in FPN-B is robust. (C) The Visual Oddball Effect map is plotted on a z scale, with the color bar at the bottom. Note further the more restricted response in the inferior parietal lobule as compared to the Old-New Recognition Effect map. (D) The group-level means of the Visual Oddball Effect estimates are plotted. (E) The Old Recognition Effect map is plotted on a z scale, with the color bar at the bottom. (F) The group-level means of the Old Recognition Effect estimates are plotted. (G) The New Recognition Effect map is plotted on a z scale, with the color bar at the bottom. (H) The group-level means of the New Recognition Effect estimates are plotted. Error bars represent the standard error of the mean across participants. * p < .05; ** p < .01; *** p < .001.

Examining the group-level maps reveals further details. The distributed regions of FPN-B are all active in the Old-New Recognition Effect and Old Recognition Effect contrasts (Figure 13A, E), including prominently the FPN-B region within IPL. This observation may explain why IPL responses were so extensive in the prior analyses extending beyond the hypothesis-targeted networks. FPN-A is also variably active, likely due to differences in controlled processing demands across the task contrasts. FPN-A’s pattern of response, like CG-OP, appears preferential to when there is an asymmetry in target status (Figure 13D, F, H). Specifically, FPN-A is active to new relative to old words in the New Recognition Effect (Figure 13G, H). By contrast, only FPN-B has the nominal properties consistent with a familiarity response^6^.

To clarify the spatial relationship between FPN-B and the observed correlates of a familiarity signal, we visualized data from the Old-New recognition Effect on a flattened cortical surface (see Figure 16 in Du et al., 2024 for correspondence between flattened and inflated views). For each participant, the Old-New Recognition Effect was overlaid with each individual’s FPN-B boundary (black outline) in the right hemisphere (Figure 14). Although the precise pattern varies across individuals, the consistent, network-wide topography suggests that the familiarity signal is distributed across FPN-B. Specifically, across multiple participants and multiple regions within each participant, the response falls clearly within the boundaries of FPN-B including a highly consistent response in IPL. The overlap extends to multiple prefrontal regions and within temporal association regions. Even within participants who show modest response, the most robust responses fall within FPN-B regions (e.g. S3 in Figure 114). Comparable organization in the left hemisphere (see Supplementary Materials) further points to a bilateral involvement of FPN-B in familiarity-related processing. All contrasts for all participants are shown in the Supplementary Materials.

**Figure 14.**
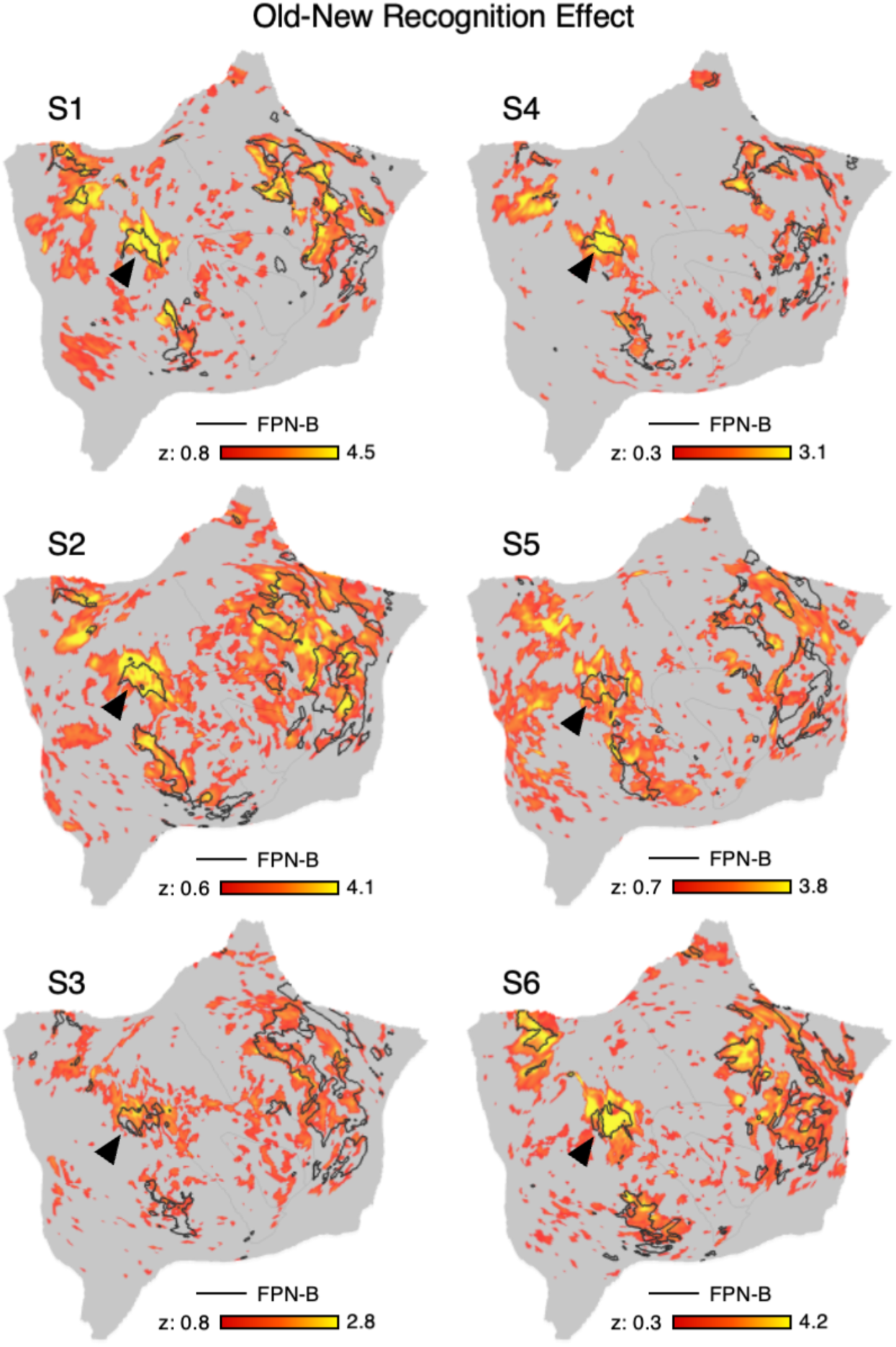
Within-individual Old-New Recognition Effect maps on the flattened cortical surface emphasize Frontoparietal Network B (FPN-B) response to familiarity. For each of six participants, the Old-New Recognition Effect is shown on a flattened cortical surface with individual FPN-B boundaries (black) overlaid on top. The maps are plotted on a z scale, with the color bar at the bottom. The format is adapted from Du et al. (2024). Across participants, the effect spans much of FPN-B, including canonical regions of FPN-B such as the inferior parietal lobule (arrows), lateral temporal cortex, and multiple regions within prefrontal cortex. Participant S3 displays among the weakest response yet the participation of the multiple regions of FPN-B is still evident including the key region of the inferior parietal lobule. This pattern is consistent with FPN-B engagement in familiarity-based processing. The Supplemental Materials show maps for both hemispheres for all participants.

## Discussion

The present study sought to disentangle component processes contributing to old-new recognition memory. We focused on a brain network, SAL/PMN, that has confounded the literature due to its emergence both as a network responding to salient stimuli across diverse contexts as well as one that responds to item repetitions in memory paradigms.

Our findings demonstrate that SAL/PMN responds robustly to salient, task-relevant events regardless of their mnemonic history. In typical memory recognition paradigms, old items may be more salient targets than new items, potentially driving responses in SAL/PMN. This conclusion was directly supported by a critical observation: SAL/PMN responded when new words were uncommon and treated as targets against a background of old words.

The network’s engagement appears to be driven by salience rather than by familiarity per se. We discuss these findings alongside additional observations that suggest another network, FPN-B, may respond to mnemonic familiarity. Thus, our results refute one hypothesis regarding a brain network supporting mnemonic familiarity, while simultaneously raising a new hypothesis to be explored in future investigations.

### SAL/PMN Responds to Salient Transient Events Not Mnemonic Familiarity

The SAL network has been proposed to contribute to detection and integration of behaviorally relevant, salient events (Seeley et al., 2007; Menon & Uddin, 2010; Seeley, 2019), whereas the PMN has been linked to familiarity-based recognition memory (Gilmore, Nelson, & McDermott, 2015; Gilmore, Kalinowski, et al., 2019). The present study was inspired by growing evidence indicating that the SAL and PMN networks are the same network studied with different emphases in parallel experimental lineages (Du et al., 2024; Kwon et al., 2025). Here we asked how salience and familiarity might interact during memory recognition tasks. Multiple observations emerged.

First, SAL/PMN could be reliably identified in each individual and be anatomically distinguished from the default network, a network juxtaposed along the posterior midline, as well as the CG-OP network, a separate network that displays multiple side-by-side adjacencies with SAL/PMN throughout the cortex (Figures 5 and 6). Second, SAL/PMN responded to both the detection of uncommon visual oddballs and the recognition of familiar old words in a standard old-new recognition task, replicating prior findings. The response was modest during old-new recognition but was present in all participants to varied degrees and robust in several. The group-average map of the Old-New Recognition Effect revealed PCU and MCC regions that are canonical to the familiarity response in the literature (e.g., compare the present Figure 8A to Figure 2 of Kim et al., 2013 and Figure 2B of Koslov, Kable, & Foster, 2024).

Third, when old words were uncommon and designated as targets, SAL/PMN exhibited a strong response, consistent with both salience and familiarity effects. Critically, when new words were uncommon and treated as targets, SAL/PMN still responded, indicating that the network’s engagement can be driven by salience even when salience is defined as novelty against a backdrop of familiar items. Thus, the response pattern can be reversed – from responding to familiar items to responding to novel ones – by altering which target is made salient. This result is inconsistent with a mnemonic familiarity hypothesis, which predicts that PMN regions should preferentially respond to familiar (repeated) items. Rather, our findings support a salience-driven account: SAL/PMN is engaged by the behavioral salience of the stimulus, not specifically its mnemonic familiarity. These findings converge with, but also clarify, earlier probability manipulations in recognition memory. In group-averaged maps, the PCU and MCC have been reported to show a familiarity (oldness) effect that appears stable across decision-criterion manipulations (Herron et al., 2004). However, group averaging and spatial smoothing can blur signals from interdigitated networks along the posterior midline (e.g., DN-A, SAL, FPN-B), obscuring network-specific responses. Using within-individual analyses, we maximized the ability to disentangle salience-related activity in SAL/PMN from contributions of neighboring networks, including FPN-B which be a source of the familiarity (oldness) effect. While being open to the possibility of additional ways in which task context may ingluence results, we are struck by the multiple observations that suggest the SAL/PMN does not respond preferentially to familiarity.

As a practical consequence of these results, we will refer to the network going forward simply as the SAL network, not SAL/PMN. A major impetus for naming the network the “Parietal Memory Network” was the finding that it responds to mnemonic repetitions (Gilmore, Nelson, & McDermott, 2015). While true under certain circumstances, the present finding that the network responds to new items under the right conditions and also responds robustly during oddball detection in non-mnemonic tasks leads us to now refer to the network solely as the SAL network, deemphasizing an implied preferential role in memory processing (see also Angeli et al., 2025). The historical name – the Salience network -- reasonably captures the current set of functional properties^7^.

### Additional Anatomical Features of the SAL Network

In addition to the finding that, under certain conditions, the SAL network responds to repeated items in memory paradigms, the SAL network is also functionally coupled to the posterior tail of hippocampus (Zheng et al., 2021). Given the critical role of the hippocampus (and related diencephalic structures) in declarative memory, the finding that the SAL network is linked to the hippocampus is also relevant to functional hypotheses. Specifically, Zheng et al. (2021) proposed that the network might participate in attention-directed memory retrieval. We previously replicated their observation that the SAL network is coupled to the posterior hippocampal tail (Angeli et al., 2025). However, we did not find response properties consistent with a traditional mnemonic role.

Rather, when examined within individuals, the posterior hippocampal region responded to oddball transients just like the SAL network itself, and further responded to task transitions between blocks of active and passive tasks that placed no explicit demands on long-term memory. Furthermore, the posterior hippocampal region linked to the SAL network could be functionally double dissociated from an anterior hippocampal region that responded during remembering and tracked the process of scene construction (Angeli et al., 2025). Thus, the response properties of the posterior hippocampal region associated with the SAL network is also consistent with a role in salience (or novelty detection) and distinct from more anterior regions.

There is another recent anatomical observation about the SAL network that deserves mention. The SAL network is also functionally coupled to the ventral striatum at or near the nucleus accumbens core extending into the ventral portion of the caudate (Lynch et al., 2024). We recently replicated and triplicated this key anatomical observation in extensive independent data (Kosakowski et al., 2025). These findings from human neuroimaging data are consistent with anatomical studies in non-human primates and rodents that have long noted direct projections between the hippocampal formation and the ventral striatum (e.g., Kelley & Domesick, 1982; Groenewegen et al., 1987; Friedman, Aggleton & Saunders, 2002; for discussion see Pennartz et al., 2011; van der Meer et al., 2014).

Taken together, a complex picture of the SAL network emerges as a brain-wide distributed network that both interacts with the posterior tail of the human hippocampus and also the ventral striatum. We do not know the functional implications of these anatomical features, and we do not yet have a clear understanding of how these anatomical observations in rodents and non-human primates relate to the human networks observed here with functional connectivity. Nonetheless, it is intriguing that there is a brain-wide distributed network that connects all these structures and responds transiently to salient behaviorally relevant events including, but extending beyond, old-new memory recognition paradigms.

### Similarities and Differences Between the SAL and CG-OP Networks

There are two separate adjacent networks that respond to salient, transient events with related but distinct properties (Seeley, 2019; Dosenbach, Raichle, & Gordan, 2025). The juxtaposed networks are referred to as the SAL network and the CG-OP network (recently renamed the Action-Mode Network, AMN; Dosenbach, Raichle, & Gordan, 2025). While they initially appeared anatomically similar to one another in group-averaged descriptions (e.g., Dosenbach et al., 2007; Seeley et al., 2007; for discussion see Seeley, 2019), precision neuroimaging investigations have demonstrated they are distinct networks with side-by-side adjacencies in multiple zones of the cortex (e.g., Gordon et al., 2017; Du et al., 2024) and in the striatum (Kosakowski et al., 2025). The present findings bolter evidence that that the two networks are similarly organized but possess distinct features. Specifically, the SAL network contains prominent regions along the posterior midline (PCU and MCC) while CG-OP does not. Rather, the CG-OP network is associated with regions that prominently surround the somatomotor cortex along the midline (Figures 5 and 6).

The two separate networks responded similarly to salient transients in our task contrasts with one key difference. Both the SAL and CG-OP networks responded during the visual oddball task (Figure 8B) but only the SAL network responded during the Old-New Recognition Effect (Figure 8A). Contrasting the SAL and CG-OP networks was not a planned comparison, but the CG-OP network shows a (slight) negative response and is entirely absent from the Old-New Recognition Effect map, as contrast to its robust, spatially specific presence in other contrasts suggesting the absence of the response may be functionally meaningful (Figure 8C, Figure 11A, C). It is thus interesting to note that all contrasts except the Old-New Recognition Effect involved a differential motor action. In the Old-New Recognition Effect, a motor response was required for both the target trials (old items) and reference trials (new items). Effects associated with motor planning and execution were presumably removed (as also evidenced by the absence of a response in left motor cortex near the hand representation). All other comparisons required a motor response to the targets while the reference trials did not.

For these reasons we speculate that, within the paradigms explored here, the CG-OP and SAL networks respond similarly with the major distinction being that the CG-OP network is linked to the motor response, consistent with recent ideas put forth by Dosenbach and colleagues that CG-OP participates, in some way, with planning motor actions (Gordon et al., 2023; Dosenbach, Raichle, & Gordon, 2025).

### A Candidate Network Responding to Mnemonic Familiarity

An unexpected finding emerged when we examined the full set of task contrasts: a right-lateralized network, FPN-B, responded consistently to attended familiar (repeated) items, but not in contrasts where salient targets lacked a familiarity signal (Figure 13). This is an intriguing result, as the functional role of FPN-B has remained somewhat enigmatic.

In Du et al. (2024), after carefully examining the task maps, we specifically noted uncertainty regarding whether any of the targeted task contrasts directly activated this network^8^. The challenge lies in FPN-B’s proximity to another network, FPN-A, which is well established in the literature as responding in a domain-flexible manner during demanding (effortful) cognitive tasks – often referred to as the multiple-demand network (e.g., Duncan & Owen, 2000; Fedorenko, Duncan, & Kanwisher, 2013; see also DiNicola, Sun, & Buckner, 2023). On a cursory inspection, FPN-B can appear to exhibit an attenuated response similar to FPN-A, likely due to spatial blur arising from their close anatomical juxtaposition. However, within-individual precision analyses – particularly in regions where the two networks are most spatially separable, such as in posterior parietal cortex – reveal that FPN-B is minimally activated by typical cognitive control demands that robustly engage FPN-A (see Figure 35 in Du et al., 2024). To date, we are unaware of any task manipulation that preferentially activates FPN-B.

Our exploratory analyses revealed that FPN-B is responsive in task contrasts that isolate the retrieval of familiar information (Figure 13). Across the four task contrasts, FPN-B was activated in a spatially preferential only when participants identified familiar, previously studied words (i.e., the Old-New Recognition and Old Recognition tasks). In contrast, FPN-B was not engaged during the Visual Oddball or New Recognition tasks, indicating that it does not track stimulus rarity or perceptual surprise. Instead, FPN-B appears to be recruited when retrieval of mnemonic information is used to guide behavior. Response maps further indicated that key parietal and prefrontal regions of FPN-B were activated, albeit not selectively. This pattern suggests a potential functional role of FPN-B in processing mnemonic familiarity. However, this interpretation remains preliminary. A comprehensive evaluation of alternative functional roles is necessary to avoid prematurely attributing a narrowly defined function to FPN-B.

## Acknowledgements

We thank the Harvard Center for Brain Science neuroimaging core and FAS Division of Research Computing. We thank T. O’Keefe for assisting in optimization of data processing and R. Mair for MRI physics support. The multi-band EPI sequence was provided by the Center for Magnetic Resonance Research (CMRR) at the University of Minnesota.

## Data Availability Statement

The data that support the findings of this study are available from the corresponding author upon reasonable request.

## IRB Statement

All participants provided informed consent using protocols approved by the Institutional Review Board at Harvard University.

## Footnotes

1 Various theoretical frameworks have been put forth, and experimental approaches adopted, to test for the presence and strength of mnemonic ‘familiarity’ signals in isolation and in the context of hypothesized recollective processes (e.g., Yonelinas, 2002; Wixted, 2007). In extreme cases, such as studied in the present work, repeated study of items leads to a strong familiarity signal that is sufficient for high performance. The task does not demand or encourage information about the study source or ancillary episodic details. Near ceiling recognition performance and tight, fast response distributions result.

2 Here we are referring to the old-new recognition tasks where mixed lists of items are presented sequentially, and the participant’s task is to quickly and accurately identify if the items are old or new. The complexity of the task increases when alternative forms of recognition decisions are used including responses that indicate confidence (Green & Swets, 1966), remember / know judgements (Tulving, 1985), application of process dissociation exclusion criteria (Jacoby, 1991), or recognition decisions that require source discrimination (Johnson & Raye, 1981). In this paper, we focus on the brain networks responding during the basic case of old-new recognition without additional decision criteria.

3 When brain responses to old items in recognition tasks were Hirst identified in fMRI neuroimaging (e.g., Henson et al., 1999; Konishi et al., 2000), parietal cortex was active leading to questions of whether attention or motor intentions were in some way confounded. As a result, the asymmetrical properties of old-new recognition tasks were noted and paradigms adopted that varied response instructions to separately target old or new items (e.g., Shannon & Buckner, 2004). We revisit these paradigms in this paper.

4 Overlapping regions are often given different names across papers. Here we refer to the PMN region along the cingulate as “mid-cingulate cortex” or MCC; the region in our precision within-network analyses extends from dorsal posterior cingulate to the mid-cingulate. The region as observed in Kim et al. (2013) is robust in the posterior cingulate and extends minimally into mid-cingulate and is labelled PCC (see Figure 1 in Kim et al., 2013). We suspect these are the same regions with slight topographic differences leading to use of different names.

5 The portion of the anterior hippocampus that is coupled to the default network, specifically default network-A (DN-A), reproducibly responds during autobiographical remembering tasks (see Figure 2 in Angeli et al., 2025). The posterior portion of the hippocampus linked to SAL/PMN responds minimally to autobiographical remembering but robustly responds to oddball transients, forming a functional double dissociation with the anterior hippocampus (Angeli et al., 2025).

6 One subtle feature of FPN-B’s response pattern is not predicted by a familiarity hypothesis. While the New Recognition Effect showed a positive response in FPN-B, it might also be expected that the response would be negative – meaning more active for the to-be-ignored old (familiar) words than the responded to new words. This was not observed. There are complexities to these tasks given their asymmetrical response demands, but in raising FPN-B’s possible involvement in mnemonic familiarity, this detail should not be ignored.

7 We generally prefer to adopt network names that define networks based on their anatomical organization since our functional understanding evolves over time (see Uddin, Yeo, & Spreng, 2019). However, many of the networks have a long history in the literature and there is generally accepted usage. Thus, we refer to the network as the Salience Network (SAL), reflecting its history and that this functional name reasonably well captures its functional response properties as surveyed to date.

8 We have been actively trying to find task contrasts that preferentially elicit a response in FPN-B for many years. We initially explored variations of cognitive control paradigms following the work of Nee and colleagues (2016; 2017) but did not obtain a preferential response, although higher-order demands on control elicited responses that overlapped and extended beyond FPN-B. We then entertained the hypothesis that FPN-B, which is right lateralized, might preferentially process non-verbal materials, but that possibility also did not pan out (e.g., Du et al., 2024). Here, in the present study targeting familiarity, we unexpectedly elicited a strong response in FPN-B.

